# From buds to shoots: Insights into grapevine development from the Witch’s Broom bud sport

**DOI:** 10.1101/2023.09.25.559343

**Authors:** Eleanore J. Ritter, Peter Cousins, Michelle Quigley, Aidan Kile, Sunil K. Kenchanmane Raju, Daniel H. Chitwood, Chad Niederhuth

**Affiliations:** Department of Plant Biology, Michigan State University, East Lansing, MI, USA; E. & J. Gallo Winery, Modesto, CA, USA; Department of Horticulture, Michigan State University, East Lansing, MI, USA; Center for Quantitative Imaging, Institute of Energy and the Environment, Penn State University; Center for Genomics and Systems Biology, New York University, Manhattan, NY, USA; Department of Computational Mathematics, Science & Engineering, Michigan State University, East Lansing, MI, USA

**Keywords:** development, grapevine, *Vitis vinifera*, liana, bud sport, somatic mutations, clonal propagation

## Abstract

**Premise of Study:** Development is relatively understudied in woody vines, such as grapevine (*Vitis vinifera*). We used the Witch’s Broom bud sport in grapevine to understand the developmental trajectories of the bud sports, as well as the potential genetic basis of the bud sport.

**Methods:** We phenotyped shoots, buds, and leaves of two independent cases of the Witch’s Broom bud sport, in the Dakapo and Merlot varieties of grapevine, alongside wild-type counterparts of the same variety. We also performed Illumina and Oxford Nanopore sequencing on the two independent cases and two wild-type counterparts of the same variety.

**Key Results:** The Dakapo and Merlot cases of Witch’s Broom displayed severe developmental defects, with no fruit/clusters formed and dwarf vegetative features. However, the Dakapo and Merlot cases of Witch’s Broom studied were also phenotypically different from one another, with distinct differences in bud and leaf development. We were able to identify unique genetic mutations in our two Witch’s Broom cases that are strong candidates for the genetic basis of the bud sports.

**Conclusions:** The Witch’s Broom bud sport in grapevine serves as a useful natural mutant in which to study grapevine development. The Witch’s Broom bud sports in both varieties studied had dwarf phenotypes, but the two instances studied were also vastly different from one another. Future work on Witch’s Broom bud sports in grapevine could provide more insight into development and the genetic pathways involved in grapevine.

## INTRODUCTION

Bud sports arise when a part of a plant, such as a lateral shoot, develops phenotypic differences from the rest of the parental plant. They typically arise when a somatic mutation occurs within a developing meristem and then spreads throughout the meristem and developing tissue (Foster and Aranzana, 2018). Bud sports are known to arise sporadically in many perennial crops and can be an important source of novel phenotypes, having given rise to many plant cultivars widely grown today. They can be an especially important source of variation in difficult to breed perennial crops, such as grapevine (*Vitis vinifera*), which is challenging to breed due to high genetic heterozygosity and long regeneration times. As a result, beneficial bud sports in grapevines have often propagated to be grown as new varieties. For example, the variety Tempranillo Blanco first arose as a bud sport of Tempranillo Tinto and was clonally propagated to maintain its novel phenotype (Carbonell-Bejerano et al., 2017). Bud sports are not always beneficial and sometimes detrimental to agricultural production, however, such bud sports provide natural mutants that can still be leveraged to study developmental traits that might otherwise not be possible (Foster and Aranzana, 2018).

Grapevines have unique development and physiology compared to many other crops and model systems. They are perennial plants that grow as lianas (also known as woody vines). This growth habit is enabled by tendrils, which are uncommon structures that allow them to climb as they grow. In addition, unlike many plants, their shoot tip does not terminate in an inflorescence, but instead contains an uncommitted primordium that allows the plants to continue growing from the tip (Gerrath et al., 2015). Development within the buds of grapevines themselves is uniquely organized to ensure the successful production of leaves, tendrils, and inflorescences from the primordia. While tendril origin differs on a species basis, grapevine tendrils are modified inflorescences (Gerrath et al., 2015). The switch from inflorescence development to tendril development occurs within the developing buds and is tightly regulated by a mixture of environmental conditions and hormones. Cytokinin signaling, high light, and high temperature promote inflorescence development while gibberellic acid (GA) signaling, low light, and low temperature promote tendril development (Srinivasan and Mullins, 1981). Changes in hormones regulating these structures can have significant impacts on the ability of *V. vinifera* to sexually reproduce, even causing seed abortion (Cheng et al., 2015). However, understanding the regulatory and genetic components involved in grapevine development has proved challenging due to the difficulty of conducting genetic and molecular studies in grapevine.

Witch’s Broom (WB) is a bud sport that occurs spontaneously in multiple grapevine varieties. The WB phenotype involves prolific vegetative growth and limited to no production of flowers (Bettiga et al., 2013). In contrast to wild-type (WT), the WB bud sport does not easily root from cuttings, although the WB sport may be propagated by grafting. Similar WB bud sport phenotypes in other plant species are usually the result of pathogen infection, typically by phytoplasma (Montano et al., 2001; Jung, 2002; Khan et al., 2002). However, genetic mutations have been shown to cause WB, as with the WB shoots in *Pinus sibirica* <u>(Zhuk et al. 2015)</u>. Cases of WB in grapevine are thought to have arisen through genetic causes and not pathogen infection. Instances of WB in grapevine do not spread within or between plants and have also occurred in plants that tested negative for pathogens. Therefore, WB bud sports in grapevines are thought to have arisen from genetic causes. As a result, the WB bud sport could be valuable for research, providing insight into an aspect of grapevine development and the genetic factors behind it, that would otherwise be near impossible to study. Here, we investigate both the phenotypic effects and the potential genetic underpinnings of two independent cases of the grapevine WB bud sport. Our results demonstrate that the WB bud sport impacts grapevine development from buds to shoots, but in distinct ways in the two cases we studied. Our work also suggests that the basis for the WB bud sports may result from mutations in different genes.

## MATERIALS AND METHODS

### Plant material***—***

Two independent cases of WB from two grapevine varieties, Merlot and Dakapo (*Vitis vinifera* L.), were sequenced and phenotyped alongside tissue from WT branches. The Merlot WT and WB samples were derived from the same plant, while the Dakapo WT and WB were derived from two separate plants.

The Merlot WB was identified as a bud sport on a vine in a commercial vineyard in Madera, California, USA that was planted in 1994. The vineyard is trained to bilateral cordons, spur pruned, and planted with rows on an east/west orientation. The proband vine was observed in 2013 to have one arm with wild type shoots (the western arm) and one arm with WB shoots (the eastern arm). The plant material both collected and studied come from a mixture of the original proband Merlot vine and cuttings derived from it. The tissue samples used for short read sequencing were collected from the contrasting arms of the original proband Merlot vine for both Merlot WT and Merlot WB. Observations and tissue samples used for long read sequencing of the Merlot WB were from the WB arm of the original proband vine as well. In 2020, budwood was collected from the WB arm of the proband vine and grafted to Rupestris St. George rootstock by a commercial nursery. The Merlot WB cuttings used for imaging buds were collected (February 2021) from those grafted Merlot WB vines planted in 2020. Cuttings from shoots on the Merlot WT arm of the proband vine were made in 2018 and the vines resulting from those cuttings were planted in 2018 and trained to bilateral cordons and spur pruned. Observations, tissue samples for long read sequencing, and cuttings of Merlot WT were collected from these planted cuttings from the proband vine.

The Dakapo WB was identified as a whole vine sport on a vine in a budwood increase block in Madera, California, USA that was planted in 2011. A budwood increase block is cultivated to provide propagation wood for grafting or cuttings rather than fruit for commercial production. The proband vine was observed in 2013 to demonstrate the WB phenotype, in contrast to nearby Dakapo WT vines of the same age in the same block. Budwood was collected from the proband vine in 2015 and grafted onto 140 Ruggeri rootstock by a commercial nursery; the grafted vines were planted in 2015. Observations and all samples of the Dakapo WB come from a single grafted vine. Observations and all samples of Dakapo WT are from the original Dakapo vines planted in the budwood increase block in 2011.

### Phenotyping of the WB bud sport***—***

Shoot and leaf phenotyping was conducted on samples from field grown vines in Madera, California, USA in September 2021. Ten shoots were examined per accession (Merlot WB, Merlot WT, Dakapo WB, Dakapo WT). For WT vines, fertile (with fruit clusters) shoots from retained nodes were observed. For WB vines, shoots from retained nodes were observed. Retained nodes are nodes with dormant buds chosen by professional pruners during dormant pruning as the most likely to produce healthy shoots in an appropriate position during the subsequent growing season and ordinarily the shoots from retained are the most fruitful shoots on a grapevine. Lateral meristem presence and type was recorded for 16 nodes beginning at the basal end of the shoot. The lateral meristem choices were tendril, cluster, and shoot. If a scar was present indicating the loss of the lateral meristem, this was recorded as “scar” since the type of lateral meristem could not be determined by observation. Skipped nodes where no lateral meristem was present were recorded as a “skip”. The length of 16 internodes basal to those nodes was recorded. The maximum blade length, maximum blade width and the petiole length of five fully expanded undamaged leaves at or distal to the cluster zone were recorded from each of the ten shoots per accession. A Welch two sample t-test was used to test for statistically significant differences between samples.

### Leaf landmarking and analysis***—***

Between 12-14 leaves were collected from six shoots per sample from plants in Madera, California, USA in June 2022. The sampled shoots grew from retained nodes. Leaves were pressed in an herbarium press at Madera, California, USA and shipped in the press to East Lansing, Michigan, USA for scanning and analysis. The leaves were scanned using a Canon CanoScan 9000F Mark II (Canon U.S.A., Inc, Melville, New York, USA) at 600 DPI. The leaves were landmarked manually by placing 21 landmarks from Bryson et al. (2020) on leaf scans using ImageJ (version 1.53k; Abramoff et al., 2004). Scans were saved as x-and y-coordinates in centimeters. The shoelace algorithm, originally described by Meister (1769), was used to calculate leaf, vein, and blade areas using the landmarks. The landmarks were used as the vertices of polygons and the following formula, as described in Chitwood et al. (2021), was used to then calculate the areas (where *n* represents the number of polygon vertices defined by the landmarked *x* and *y* coordinates):

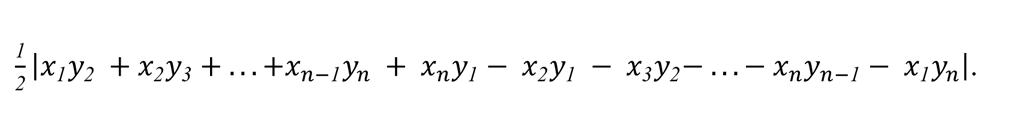

A Welch two sample t-test was used to test for differences in leaf area between samples. To investigate changes in leaf shape between WT and WB leaves, a generalized Procrustes analysis and a principal components analysis (PCA) was performed using the shapes package (version 1.2.7; Dryden and Mardia, 2016) in R (version 4.2.2; R Core Team, 2022) and RStudio (version 2022.12.0.353; RStudio Team, 2022), with scaling and rotation. The shapes package (version 1.2.7; Dryden and Mardia, 2016) in R and RStudio was also used to test for mean shape differences using a Hotelling’s *T^2^*test.

### Data visualization***—***

All plots were made in R using ggplot2 (version 3.4.2; Wickham, 2016) and arranged using cowplot (version 1.1.1; Wilke, 2020). The R package ggsignif was used to add significance bars to violin plots (version 0.6.4; Constantin and Patil, 2021). The R package ggnewscale was used to plot distinct scales for WT and WB data when needed (version 0.4.8; Campitelli, 2023).

### Bud collection, dissecting, and imaging***—***

Dormant grapevine cuttings were collected in Madera, California, USA in February 2021 and shipped overnight to East Lansing, Michigan, USA. The Dakapo WT and Merlot WT cuttings were between 6-7 mm in diameter, while the Dakapo WB and Merlot WB cuttings were between 4-5 mm in diameter. Cuttings were left at room temperature for 24-72 hours before dissecting. Only live cuttings were used for bud dissection. The buds were dissected using a razor, slicing the buds vertically (parallel to the stem) until the primary, secondary, and tertiary buds could all be seen, but tendril primordia were still distinguishable. Buds were then imaged with a dissecting microscope.

Buds were also scanned to create 3D X-ray Computed Tomography (CT) reconstructions of internal anatomy. The scans were produced using the North Star Imaging X3000 system and the included efX software (Rogers, Minnesota, USA). The scans were taken at 75 kV and 100 µamps with a frame rate of 12.5 frames per second in continuous mode. 2880 projections and 2 frame averages were used. To obtain the maximum voxel size (4.5 µm), a subpix scan, which takes 4 scans at half a pixel distance and combines them to get approximately half the voxel size, was used (see scale, Fig. 5). The 3D reconstruction of the buds was computed with the efX-CT software. efX-View software was used to visualize 2D slices through the 3D reconstructions of the buds.

### Whole genome sequencing and alignment***—***

Leaf tissue samples for sequencing were collected from all four accessions (Merlot WB, Merlot WT, Dakapo WB, Dakapo WT) in August 2018. DNA isolation was performed using the CTAB method as described in (Porebski et al., 1997). Library preparation for paired-end (PE) sequencing was performed as in (Urich et al., 2015) with slight modification and sequenced on an Illumina HiSeq 2500 (Illumina, Inc., San Diego, California, USA) with 150 base pair (bp) PE reads sequenced to 50-58X coverage. The reads were then prepared for downstream analysis, first using cutadapt (version 3.7; Martin, 2011) to trim adapters and low-quality bases from the beginning and ends of reads. The quality of the reads, both before and after trimming, were checked using FastQC (version 0.11.9; Andrews, 2010). The trimmed reads were then mapped to the 12X.v2 grapevine reference genome assembly (Canaguier et al., 2017) using BWA-MEM and the *-M* parameter (version 0.7.17; Li and Durbin, 2009). Mapped reads were then prepared for variant calling by sorting them with Samtools (version 1.9; Danecek et al., 2021) and marking duplicate reads using Picard MarkDuplicates (version 2.15.0; Broad Institute, 2019). The reads were then indexed using Samtools (version 1.9; Li et al., 2009), to enable use with downstream variant callers.

### SNP calling and annotation***—***

The GATK (version 4.0.12.0; McKenna et al., 2010) pipeline for short variant discovery was used to call SNPs in the samples using the BAM files with marked duplicates (DePristo et al., 2011). GATK HaplotypeCaller was used to call SNPs in the individual samples. The SNPs were combined into one file and genotyped using GATK CombineGVCFs and GenotypeGVCFs, respectively. The SNPs were filtered with GATK VariantFiltration (DePristo et al., 2011; Van der Auwera et al., 2013), using the following filters: *MQ<40.00, FS>60.0, QD<2.0, MQRankSum<-12.5,* and *ReadPosRankSum<-8.0*. ANNOVAR was used to annotate the SNPs (version 2018-04-16; Wang et al., 2010) with the Genoscope 12X grapevine genome annotation (Jaillon et al., 2007) lifted to the 12X.v2 grapevine genome assembly (Canaguier et al., 2017) using liftoff (version 1.6.2; Shumate and Salzberg, 2021) to minimize compatibility issues the newest grapevine genome annotation (Canaguier et al., 2017) had with downstream analyses.

### Long read sequencing***—***

New tissue was collected for Oxford nanopore technology (ONT) sequencing in July 2021. The tissue samples used were young leaves collected from actively growing shoot tips. The samples were frozen and shipped on dry ice overnight. The MSU Genomics Core extracted DNA from the samples and prepared the sequencing libraries. DNA was isolated from samples using a modified Qiagen Genomic-tip protocol (Qiagen, 2015) with 5 mg lysing enzyme (0.5 mg/ml; Sigma Cat# L1412-5G; Sigma-Aldrich, Inc., St. Louis, Missouri, USA), 5 mg Pectinase (0.5mg/ml; Sigma Cat# P2401; Sigma-Aldrich, Inc., St. Louis, Missouri, USA), and 500 µl Viscozyme L (5%; Millipore Sigma Cat# V2010-50; MilliporeSigma, Burlington, Massachusetts, USA) added to the lysis buffer. Short read elimination was performed using the Circulomics Short Read Eliminator kit (formerly SS-100-101-01, now SKU 102-208-300; PacBio, Menlo Park, California, USA). The size selected DNA was quantified using a Qubit 1.0 Fluorometer (Thermo Fisher Scientific, Waltham, Massachusetts, USA) and the Qubit dsDNA BR (Broad Range) Assay (Q32853; Thermo Fisher Scientific, Waltham, Massachusetts, USA). Barcoded sequencing libraries were then prepared using the Oxford Nanopore Technologies Ligation Sequencing Kit 1D (SQK-LSK109) and Native Barcoding Expansion Kit (EXP-NBD104) (Oxford Nanopore Technologies, Oxford, UK). The pooled libraries were then loaded on a PromethION FLO-PRO002 (R9.4.1) flow cell and sequenced on a PromethION24, running MinKNOW Release 21.11.7, to 17-28.5X coverage (Oxford Nanopore Technologies, Oxford, UK). Base calling and demultiplexing were done using Oxford Nanopore’s Guppy (version 5.1.13; Oxford Nanopore Technologies, Oxford, UK) with the High Accuracy base calling model.

### Long read alignment and structural variant calling***—***

Adapters were trimmed from the ONT reads using Porechop (version 0.2.4; Wick et al., 2017). NanoLyse was used to remove ONT reads mapping to the lambda phage genome (version 1.2.0; De Coster et al., 2018). Low-quality reads and reads shorter than 300 base pairs (bp) were removed using NanoFilt (version 2.8.0; De Coster et al., 2018). The quality of the trimmed and filtered reads was analyzed using FastQC (version 0.11.9; Andrews, 2010).

The ONT reads were mapped to the 12X.v2 grapevine reference genome assembly (Canaguier et al., 2017) using minimap2 (version 2.23-r1111; Li, 2021) twice with different parameters based on the needs of downstream programs. For use with sniffles (version 1.0.12; Sedlazeck et al., 2018) to call structural variants (SVs), ONT reads were mapped with minimap2 and the following parameters: *-ax map-ont --MD*. The mapped reads were sorted with SAMtools (Danecek et al., 2021). Sniffles was first run on sorted mapped read files for all samples separately using the *--snf* parameter to generate .snf files for all samples. Sniffles was then run on the .snf files previously generated for WT and WB samples from the same variety, running Dakapo and Merlot separately, to create a VCF file with SVs (version 1.0.12; Sedlazeck et al., 2018).

The second version of ONT read mapping used minimap2 (version 2.23-r1111; Li, 2021) with parameters optimized for use with pbsv (version 2.8.0; Pacific Biosciences, 2021), an additional SV caller: *-a --MD --eqx -L -O 5,56 -E 4,1 -B 5 --secondary=no -z 400,50 -r 2k -Y*. Samtools (version 1.9; Danecek et al., 2021) was used to sort the mapped reads and add read groups. The sorted mapped read files were then used with pbsv “discover”, running all samples separately to first discover signatures of structural variation and produce a. svsig file. A VCF file with SVs was then generated by running pbsv “call” (version 2.8.0; Pacific Biosciences, 2021) with .svsig files for WT and WB samples from the same variety (with Dakapo and Merlot samples run separately) and the 12X.v2 grapevine reference genome assembly (Canaguier et al., 2017).

The SVs generated by sniffles and pbsv were first filtered to remove variants that did not pass the filters applied by the two variant callers. For total structural variant counts by sample, the filtered VCF files from sniffles and pbsv were then merged using SURVIVOR “merge” (version 1.0.7; Jeffares et al., 2017) to merge SVs identified by both programs that were greater than 30 bp long and within 300 bp of one another. To identify variants with genotypes specific to the WB samples and not present in WT, SnpSift (version 2017-11-24; Cingolani et al., 2012) was used with the filtered VCF files to extract out variants either a) only found in the WB sample (homozygous or heterozygous) or b) homozygous in the WB sample and heterozygous in the WT sample. The VCF files filtered both by quality and SnpSift from sniffles and pbsv were then merged using SURVIVOR “merge” (version 1.0.7; Jeffares et al., 2017) as described previously. Only SVs that met those two criteria for merging were used for downstream analysis. The genes overlapping with the merged SVs were identified using bedtools “intersect” (version 2.30.0; Quinlan and Hall, 2010) and the Genoscope 12X grapevine genome annotation (Jaillon et al., 2007) lifted to the 12X.v2 grapevine genome assembly (Canaguier et al., 2017) using liftoff (version 1.6.2; Shumate and Salzberg, 2021).

### Candidate gene analysis***—***

To investigate a potential causal gene(s)/variant(s) for the WB budsport in grapevine, all genes with high impact SNPs or SVs present in the WB samples and either a) absent in WT (described as “novel” from hereinafter) or b) heterozygous in WT but homozygous in WB (described as “genotypically distinct” from hereinafter), were investigated for gene function by looking into the functions of their closest *Arabidopsis thaliana* ortholog. In order to understand the putative functions of the genes with SNPs and SVs in the WB samples, diamond (version 0.8.36; Buchfink et al., 2015) was used to search for *Arabidopsis* orthologs to the putative causal genes using the Araport 11 *Arabidopsis* annotation (Cheng et al., 2017). The list of *Arabidopsis* genes orthologous to WB candidate genes was loaded into RStudio, and the R/Bioconductor package biomaRt (version 2.54.1; Durinck et al., 2009) was used to obtain gene descriptions from Ensembl Plants (Bolser et al., 2016). The *Arabidopsis* orthologs and the information about their function were then used to prioritize genes involved in developmental, hormone signaling, or other pathways that could potentially result in the WB phenotype. Variants of interest were verified first by looking at mapped reads for all samples in a genome browser to verify that the genetic variants were truly genotypically distinct to the WB sample. Then, polymerase chain reaction (PCR) was used to validate the variant in all samples. The amplified products were Sanger sequenced to verify that the variant called was accurate in both location and genotype.

## RESULTS

### WB shoot phenotypes***—***

The WB bud sport arises spontaneously in many varieties of grapevine (Bettiga, 2013). We characterized two independent cases of WB that occurred at a commercial vineyard in Madera, CA. The first case is a WB mutant of a Merlot grapevine, observed as one arm (the eastern) on a vine in a commercial vineyard block. The adjacent western arm on the same plant is WT. This allowed a direct comparison of WB and WT tissues from the same plant. The second case characterized was in the Dakapo variety and is a WB vine that was identified as a whole plant mutation. As a result, no WT shoots were present on the Dakapo WB plant, so separate, unaffected Dakapo vines from the same propagation batch were used as the WT comparison. In both cases, the bud sport is characterized by vigorous vegetative growth with shortened internodes (Fig. 1). Merlot WB shoots have light green leaves strikingly distinct from WT shoots (Fig. 1A), while Dakapo WB leaves are similar in color to WT shoots (Fig. 1C).

**Fig. 1.**
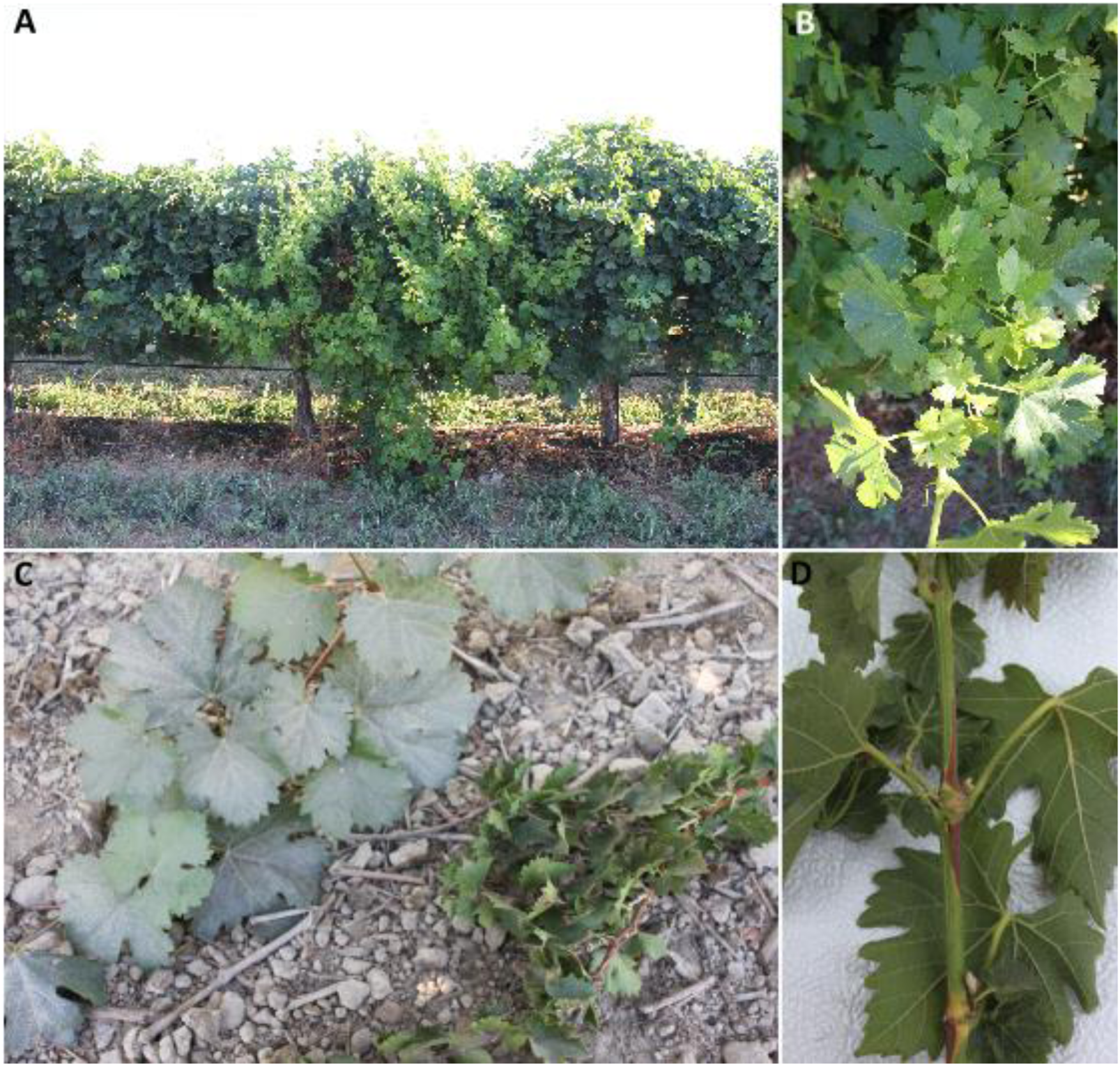
Photos of wild-type and Witch’s Broom shoots from a commercial vineyard. **(A)** Photos of Merlot WB and WT on one grapevine plant. WB shoots are the light green shoots in the center of the image, while WT shoots are the darker green shoots on either side of the WB shoots. Merlot WB shoots display prolific growth in comparison to their WT counterparts. **(B)** An up-close photo of Merlot WB shoot, with light green leaves and shortened internodes. **(C)** A side-by-side photo of Dakapo WT (left) and Dakapo WB (right) shoots from different plants. Dakapo WB shoots have shortened internodes and more prolific foliage than their WT counterparts. **(D)** An up-close photo of a Dakapo WB shoot, showing a significantly shortened internode.

Comparison of multiple shoot traits between the WT and WB plants revealed large differences in phenotypes between the two. Both Dakapo and Merlot WB shoots have internodes significantly shorter than their WT counterparts (t = -21.86, df = 230.76, P < 0.001 for Dakapo; t = -2.93, df = 317.25, P = 0.003 for Merlot) (Fig. 2A). The petioles were also smaller in WB plants than WT (t = -27.72, df = 32.91, P < 0.001 for Dakapo; t = -5.01, df = 87.44, P < 0.001 for Merlot) (Fig. 2B). Our phenotyping also revealed that the Dakapo WB phenotype seems to be more severe than the Merlot WB phenotype. The Dakapo WB internodes are significantly shorter than those of Merlot WB (t = -15.54, df = 281.57, P < 0.001), despite Dakapo WT internodes being longer than Merlot WT internodes (t = 6.58, df = 294.07, P < 0.001) (Fig. 2A). In addition, the Dakapo WB petioles are also significantly shorter than their Merlot WB counterparts (t = -25.19, df = 69.41, P < 0.001) (Fig. 2B).

**Fig. 2.**
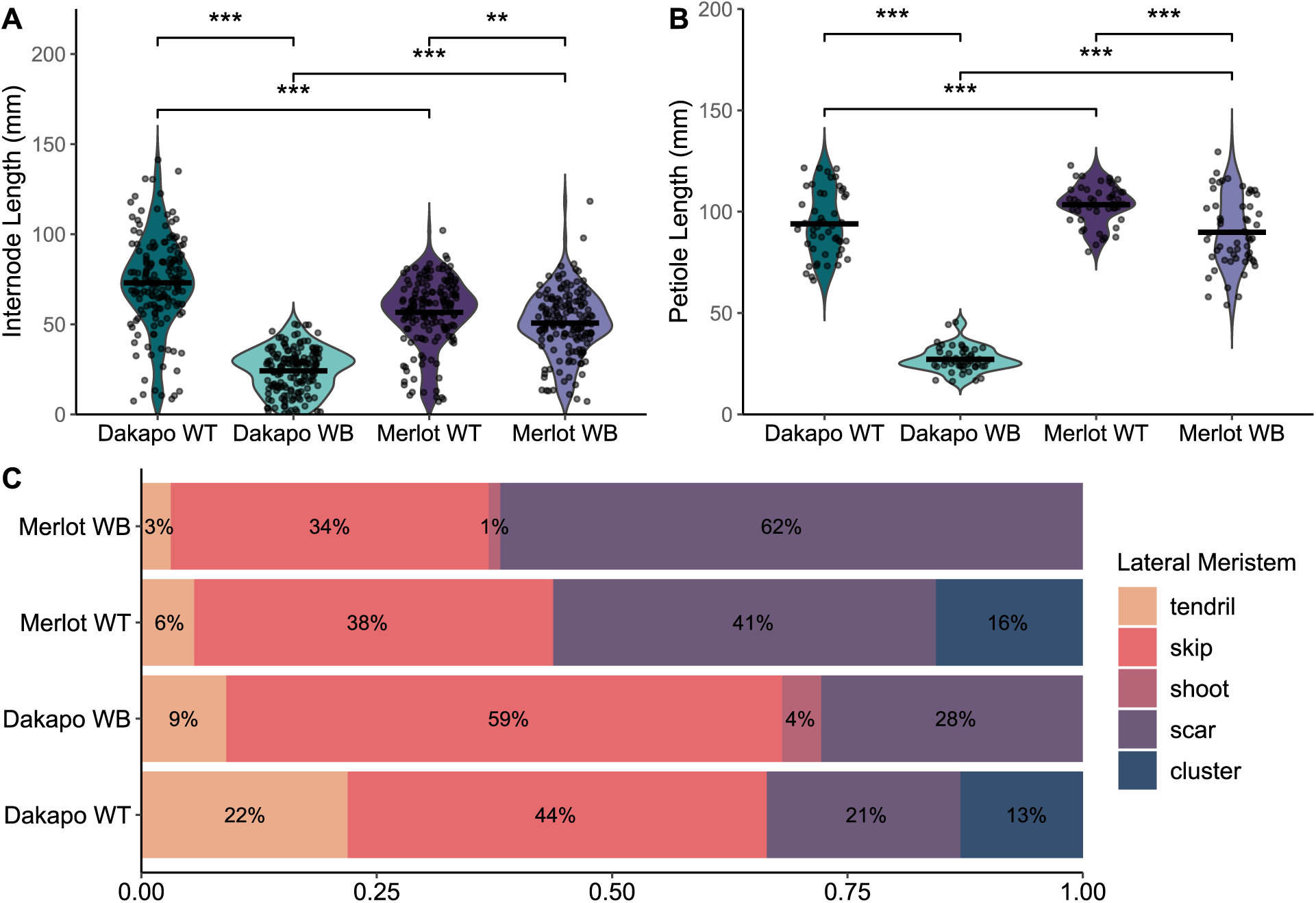
Differences in shoot phenotypes between wild-type and Witch’s Broom samples in Dakapo and Merlot varieties of grapevine. **(A)** A comparison of average internode length and **(B)** petiole length between sample types, collected from 10 shoots each. Mean values were represented by a black line for each sample. Dakapo WB and Merlot WB both have significantly smaller internodes (P<0.001*** and P<0.01**, respectively) and petioles (P<0.001*** for both cases) in comparison to WT plants of the same variety. The WT samples of the two varieties differ as well, with Dakapo WT having longer internodes but shorter petioles than Merlot WT (P<0.001*** for both). Dakapo WB also has significantly smaller internodes and petioles compared to Merlot WB (P<0.001*** for both). **(C)** The percentage of nodes with specific lateral meristem outcomes, collected from 144-160 lateral meristems for each sample.

Initial measurements of leaf width and length demonstrated that Dakapo and Merlot WB leaves are significantly shorter and narrower than their WT counterparts when compared at the same node (P < 0.05 for width and length at node 4 for Dakapo; P < 0.05 for both width and length, for nodes 5-9 for both Dakapo and Merlot cases) (Appendix S1). While initial data collected in 2021 showed that the Dakapo and Merlot WB leaves were typically shorter and narrower than their WT counterparts (Appendix S1), the actual change in leaf area and leaf shape was unknown. Leaves collected and landmarked from all samples in 2022 demonstrated that WB leaf areas were significantly smaller overall than their WT counterparts (t = 23.49, df = 76.98, P < 0.001 for Dakapo; t = 22.41, df = 70.42, P < 0.001 for Merlot) (Fig. 3A-E). To further understand how WB leaf development may differ from typical grapevine leaf development, we calculated the allometric ratio of vein area to blade area. As leaves expand, the blades of leaves expand at a greater rate than the veins (Chitwood et al., 2016). As a result, larger leaves typically have lower vein-to-blade ratios. In addition, the ratio of vein-to-blade area is typically more responsive to subtle changes in leaf shape and development than area alone (Chitwood et al., 2021). As expected, given their small leaves, both Dakapo WB and Merlot WB have significantly higher vein-to-blade ratios in comparison to WT plants of the same variety (t = -16.67, df = 133.14, P < 0.001 for Dakapo; t = - 19.08, df = 127.55, P < 0.001 for Merlot) (Fig. 3F). Dakapo WB leaves also have a higher vein-to-blade ratio than Merlot WB leaves (t = 6.53, df = 120.44, P < 0.001) (Fig. 3F). This is likely due to very subtle differences in leaf development between the two WB samples that are not captured by comparing leaf area alone, such as differences in vasculature development between the two.

**Fig. 3.**
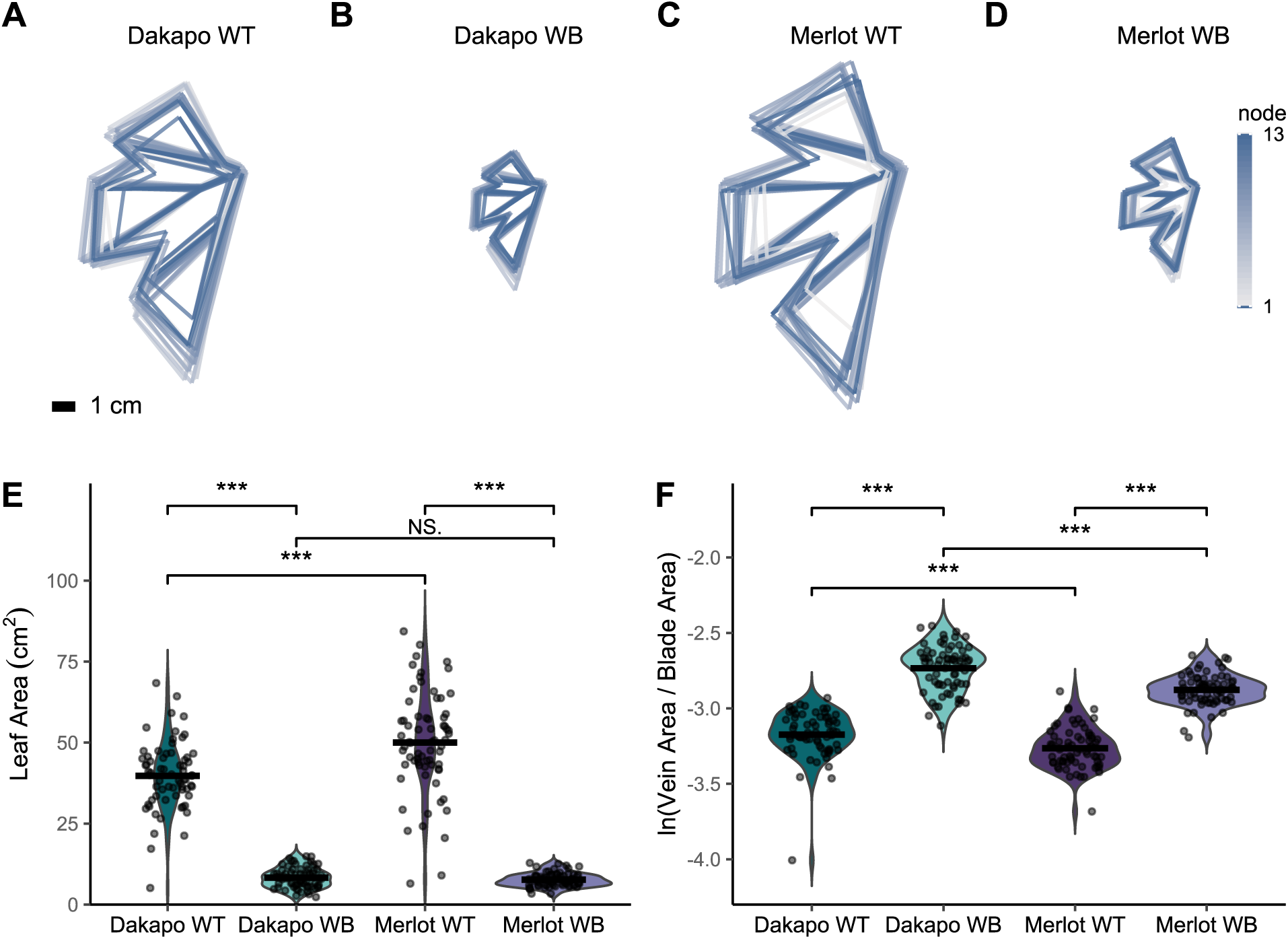
Comparing leaf area and the natural log of the ratio of vein-to-blade area between wild-type and Witch’s Broom samples in Dakapo and Merlot varieties of grapevine. **(A)** Dakapo WT, **(B)** Dakapo WB, **(C)** Merlot WT, and **(D)** Merlot WB composite leaves generated using leaf landmarks to model leaf shapes for leaves collected across 13 nodes. Composite leaves are colored based on node, from gray (node 1 from the shoot tip) to dark blue (node 13). All samples are to the same scale, and a 1 cm scale bar is provided in the bottom left corner of **(A)**. **(E)** A comparison of leaf area (cm^2^), as calculated using the shoelace algorithm originally described by Meister (1769) and used in Chitwood et al. (2020) to calculate leaf area in grapevine, with leaf landmark data. Mean leaf area (cm^2^) is represented by a black line for each sample. Dakapo WB and Merlot WB both have significantly smaller leaves (P<0.001*** for both cases) in comparison to WT plants of the same variety. Merlot WT leaves were larger than Dakapo WT leaves (P<0.001***), however leaf area did not differ between the two WB cases (P=0.16). **(F)** A comparison of the natural log of the ratio of vein-to-blade area, an allometric indicator of leaf size that is typically more sensitive to leaf size changes than leaf area alone. Mean ln (vein-to-blade ratio) is represented by a black line for each sample. Dakapo WB and Merlot WB both have significantly higher vein-to-blade ratios (P<0.001*** for both cases) in comparison to WT plants of the same variety. Dakapo WT leaves have a higher vein-to-blade ratio than Merlot WT leaves (P<0.001***). Dakapo WB leaves have a higher vein-to-blade ratio than Merlot WB leaves (P<0.001***) as well.

Analyzing the leaf landmark data utilizing a Procrustes analysis and a principal components analysis (PCA) revealed that WB leaves also differ in their shape when compared to their WT counterparts (H = 13.26, P < 0.001 for Dakapo; H = 14.07, P < 0.001 for Merlot) (Fig. 4). Eigenleaves from the PCA comparing leaf shape between scaled WT and WB leaves (Appendices S2 and S3) revealed the shape features that each PC reflected. The leaf shape variance between WT and WB in both Merlot and Dakapo appears to be due to similar phenotypic changes in the WB leaves. For both varieties, PC2 reflects variance in the depth of the distal sinus, which is deeper in WB samples. WB leaves in both varieties also seem to have a wider petiolar sinus, which is reflected by PC3 in Dakapo and PC4 in Merlot. Additionally, WB plants in both varieties appear to have narrower upper lateral lobes, which is explained by PC4 in Dakapo and PC1 in Merlot (Fig. 4). Despite these similarities in how WB leaves differ from WT in the two varieties, WB also appears to impact leaf shape somewhat differently in the two varieties. Dakapo WB leaves appear to have narrower distal sinuses than their WT counterparts, as described by PC1 (Fig. 4A, C). Meanwhile, Merlot WB leaves appear to have shorter midveins relative to the rest of leaf features, as explained by PC3 (Fig. 4B, D). These two features appear to be specific to WB bud sports of the particular variety.

**Fig. 4.**
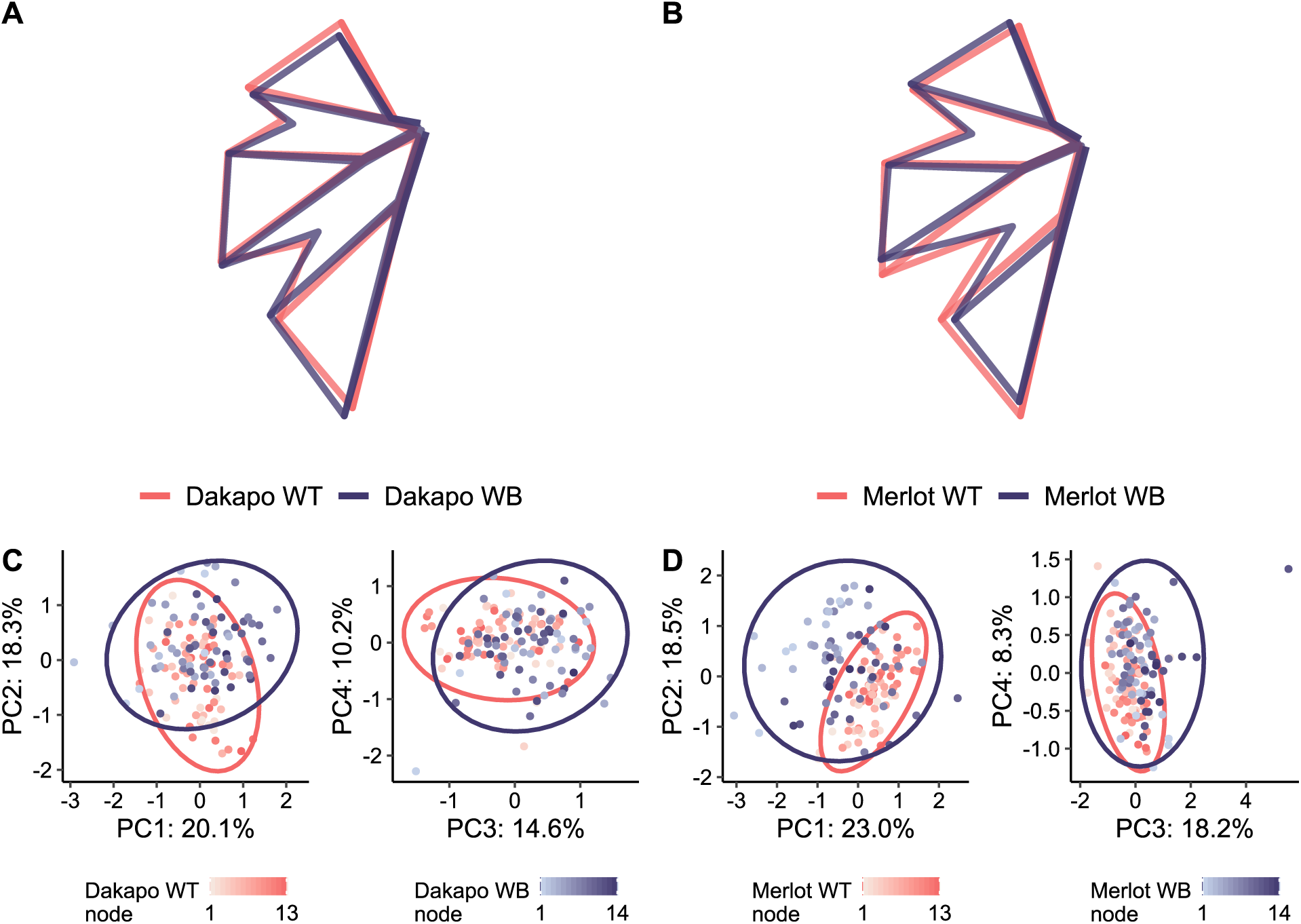
Mean leaf shapes rotated and scaled identically for **(A)** Dakapo WT and Dakapo WB, as well as for **(B)** Merlot WT and Merlot WB. **(C and D)** Principal component analysis (PCA) of all leaf shapes, with WT colored in salmon and WB colored in purple, for **(C)** Dakapo and **(D)** Merlot. The node position of the leaves is also shown by shade, with the lightest shade being node 1 (from the shoot tip) and the darkest shade being node 13-14, depending on the sample.

We also characterized the fates of lateral meristems of the WB bud sports to understand the developmental outcomes of the WB buds. Lateral meristem fates were characterized by the organ or structure that had developed at the nodes, which were either: a) tendrils, b) skips (nodes where no lateral meristem was present), c) shoots, d) scars (nodes where a meristem had formed, but no structure was present when phenotyped), or e) clusters/fruit. These observations revealed that no clusters were developing in the WB shoots. In addition, the WB shoots developed new lateral shoots at 1-4% of nodes, while their WT counterparts did not develop these new lateral shoots at any nodes (Fig. 2C). New grapevine shoots arise from axillary buds, and lateral shoots typically do not develop. It is possible that the incidence of lateral shoots on the WB bud sports may be due to the mutation directly. Both the presence of the lateral shoots and absence of clusters support that the WB bud sports seem to involve a shift towards vegetative growth and away from reproductive growth. Many of the WB lateral meristems failed to develop properly, with 87% of Dakapo WB buds and 96% of Merlot WB buds failing to develop into tendrils, clusters, or shoots, compared to 65% and 79% in their respective WT counterparts. The higher incidence of skips in Dakapo WB (59%), in comparison to Dakapo WT (44%), contributes directly to the lack of tendrils and clusters observed. Dakapo WB having more skips present is somewhat unexpected, as grapevines are expected to generally show a phyllotaxy of two successive nodes with a lateral meristem followed by one node without. As a result, we generally expect to see ⅓ of the nodes studied to be skips. It is possible that the WB mutation in Dakapo causes an unusual phyllotaxy and thus more skips to be present. However, the Merlot WB shoots have about the expected number of skips present (34%). While the characterization of lateral meristem fate demonstrated dominance of vegetative growth in both instances of WB, it also revealed that they may have distinct issues when it comes to lateral meristem development.

### Organization and development of WB buds***—***

To investigate the developmental origin and timing of the defects seen across the WB shoots, particularly in lateral meristem fates, dormant winter buds were imaged to identify changes in bud organization. To do so, we imaged dissected grapevine buds with a dissecting scope and whole buds with CT scans. Grapevine dormant winter buds are typically composed of three bud primordia, characterized as primary, secondary, and tertiary, from most developed to youngest respectively. The bud primordia typically house leaf, tendril, and inflorescence primordia (Gerrath et al., 2015). The WT buds for both Dakapo and Merlot varieties had nearly identical organization and structures. The buds and primordia were each at a 45° angle from the stem. All WT buds had three bud primordia in each of the buds as expected. CT scans showed that all WT buds had inflorescence primordia present, with 80% of WT buds having two or more inflorescence primordia present in their primary bud primordia alone. None of the WT buds appeared to have any organizational defects in the buds, with all primordia appearing to be healthy and properly arranged (Fig. 5A, C and appendix S4A, C).

In contrast, the WB buds contained multiple organizational defects. Upon examination, about half of the Dakapo WB buds had an initiated lateral shoot stem extended out of them, about 1 cm long (Appendix S4B). CT scans revealed that this stem appears to be vascular tissue pushing through the bud, disrupting the typical organization. The vascular tissue expanding through the buds sometimes contained a shoot apex on the tip, suggesting that these shoots can produce leaves and other lateral organs. Half of the buds contained an additional change in overall architecture, with the tertiary primordia being perpendicular to the stem (Fig. 5B). Many of the primordia present in the Dakapo WB buds appeared to be smaller than those in the other samples. Notably, three of four Dakapo WB buds had only one inflorescence primordia present, but the inflorescence primordia appeared deformed in two of the buds scanned.

**Fig. 5.**
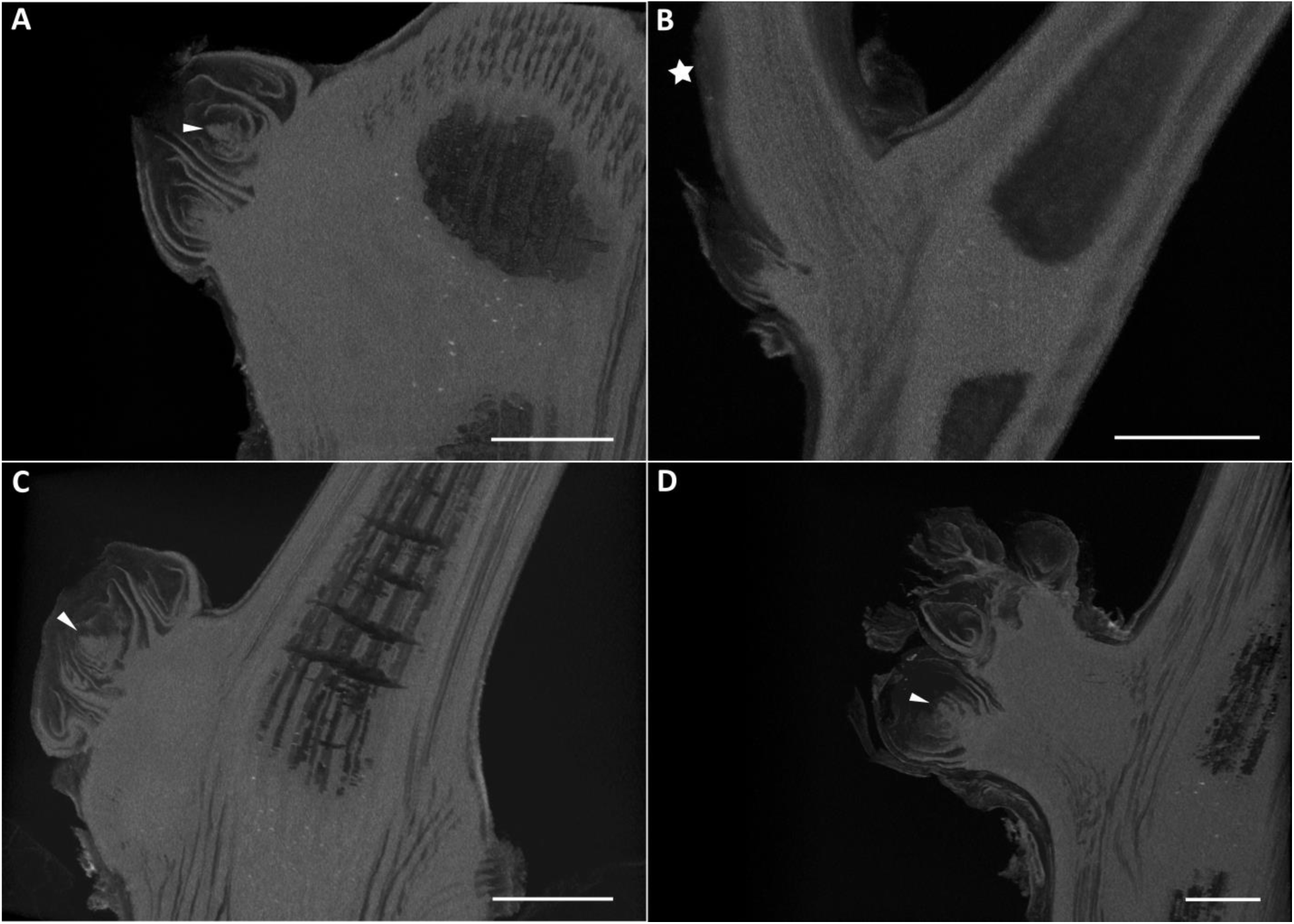
CT scans of buds from **(A)** Dakapo WT, **(B)** Dakapo WB, **(C)** Merlot WT, and **(D)** Merlot WB samples. The inflorescence primordia are indicated by the solid triangle in the **(A)** Dakapo WT, **(C)** Merlot WT, and **(D)** Merlot WB samples. The initiated lateral shoot stem in the **(B)** Dakapo WB sample is indicated by the solid star. Only one inflorescence primordium is present in the images, although multiple were seen for both WT samples. The Merlot WB sample shown **(D)** is the only Merlot WB sample scanned with a potential inflorescence primordium present, although the inflorescence primordia seen appears to be deformed. The potential Merlot WB inflorescence primordium **(D)** has smoother edges than those seen in WT samples (**A and C)**. Scale bar = 1 mm.

The Merlot WB buds had drastically different organization from WT buds as well, with the buds containing between 4-8 bud primordia (Fig. 5D and appendix S4D), in contrast to the 3 consistently found in wild-type samples (Fig. 5A, C and appendix S4A, C). Similarly to the Dakapo WB samples, two out of five of the Merlot WB buds had tertiary primordia nearly perpendicular to the stem. In addition, all but one of the Merlot WB buds had no inflorescence primordia. The inflorescence primordium potentially present in the single sample was difficult to confidently identify as such, however, since it lacks the lobes typically seen in developing inflorescence primordia (Fig. 5D). As a result, even if this structure is truly an inflorescence primordium, it is extremely deformed. However, none of the Merlot WB buds displayed the vascular tissue expansion seen in the Dakapo WB samples.

Overall, the Dakapo and Merlot WB buds contained phenotypes vastly different from WT and even one another. The WB samples displayed extensive defects in bud organization and the quantity of inflorescences produced indicating that the WB defects manifested early in bud development. This investigation into the buds of the WB bud sports provided insight into the defects we identified across the shoots of the bud sports. Not only are the shoots failing to develop properly, but the defects are pervasive in the buds and potentially their internal structures, as well.

### Genetic variation in WB Bud Sports***—***

To investigate the genetic basis of the WB bud sport, we sequenced DNA from both Dakapo and Merlot WB and WT using both Illumina 150 bp paired-end sequencing and Oxford Nanopore long-read sequencing. The average read depth coverage of the Illumina reads was between 32-37X for the samples. The average read depth coverage of the Oxford Nanopore reads was between 16-27X for the samples. The read length N50 for the trimmed Oxford Nanopore reads was between 12,890-14,486 bp for the samples. High quality reads were used for mapping to the reference genome and calling variants in each of the samples. For all samples, over 98.2% of Illumina reads and over 99.9% of the Oxford Nanopore reads mapped to the grapevine 12X.v2 reference genome (Canaguier et al., 2017).

SNPs were called against the 12X.v2 grapevine reference genome (Canaguier et al., 2017) using Illumina sequencing data. Each sample had between 7.9-8.2 million SNPs and high heterozygosity (67.96-71.22%). Most SNPs were present in both WT and WB samples of the same variety (94.81-94.97%), however between 409,588-418,818 SNPs were entirely novel when compared within-variety. A majority of SNPs were either intergenic or not expected to have an impact on gene function (Appendix S5). Of the SNPs called in the WB samples, 6,296-6,450 SNPs were predicted to have high impact on gene function. Between 597-613 genes impacted by SNPs predicted to have a high impact were genotypically distinct in WB bud sports, and these genes were kept as possible causal candidates for the bud sport (Table 1).

**Table 1.** Genetic variants identified in samples when called against the 12X.v2 grapevine reference genome assembly, including SNPs and SVs. Novel and genotypically distinct SNPs were identified by comparing variants intra-variety.

|  | **Dakapo** |  | **Merlot** |  |
| --- | --- | --- | --- | --- |
|  | **WT** | **WB** | **WT** | **WB** |
| **SNPs** |  |  |  |  |
| Total SNPs | 7,912,797 | 7,925,441 | 8,148,571 | 8,163,074 |
| Genotypically distinct SNPs<sup>a</sup> | 497,531 | 493,394 | 495,782 | 503,973 |
| Novel SNPs<sup>b</sup> | 410,608 | 411,425 | 409,588 | 418,818 |
| High impact SNPs | 14,013 | 13,962 | 14,032 | 14,027 |
| Genes impacted by high impact SNPs<sup>c</sup> | 6,310 | 6,296 | 6,450 | 6,445 |
| Genes impacted by genotypically distinct high impact SNPs | 611 | 613 | 591 | 597 |
| **SVs** |  |  |  |  |
| Total SVs | 52,119 | 53,089 | 54,775 | 53,912 |
| Genotypically distinct SVs<sup>a</sup> | 578 | 635 | 691 | 540 |
| Novel SVs<sup>b</sup> | 157 | 223 | 224 | 102 |
| Genes impacted by SVs | 15,044 | 15,134 | 15,706 | 15,596 |
| Genes impacted by genotypically distinct SVs | 135 | 136 | 150 | 134 |

<sup>a</sup> Variants genotypically distinct from the sample of the same genotype include entirely novel SNPs, as well as SNPs that have different genotypes when compared intra-variety.
<sup>b</sup> Novel variants are variants completely absent in the sample of the same variety.
<sup>c</sup> High impact SNPs include frameshift deletion or insertion, stop gain/loss, and splicing.

Structural variants were called against the 12X.v2 grapevine reference genome (Canaguier et al., 2017) using long-read sequencing data. Each sample had between 52-55,000 SVs. Deletions were the most common type of SV and accounted for over half of the SVs called. Insertions were the next most common type of SV and accounted for about 47% of total SVs. Inversions, transversions, and duplications were extremely rare, and collectively only accounted for between 1.27-1.66% of all SVs called (Appendix S6). Entirely novel SVs (when compared within variety) were rare as well, with only between 102-224 identified within the samples. Only 635 and 540 SVs were genotypically distinct in Dakapo WB and Merlot WB, respectively. About 15,000 genes had SVs within them for each sample individually. Of the genes containing SVs, 136 and 134 were impacted by genotypically distinct SVs for Dakapo WB and Merlot WB, respectively (Table 1).

We looked at the gene function for 577 genes unique to Dakapo WB and 561 genes unique to Merlot WB. The two WB samples shared 164 genes impacted by variants. All of the genes in common between the two WB samples were weak candidates with either gene functions unrelated to the WB phenotype or were unsupported by the genome browser and/or PCR validation. While we were unable to confidently identify a single, strong putative candidate gene for Dakapo WB, our variant calling pipelines identified 974 potential candidates for the causal variant(s) of the Dakapo WB bud sport (Appendix S7). In Merlot WB we identified as well as 968 potential candidates (Appendix S7), including one strong putative candidate gene, GSVIVG01008260001, an ortholog of *Arabidopsis* STOMATAL CYTOKINESIS-DEFECTIVE 1. The Merlot WB bud sport samples contain a heterozygous 3.6 kbp insertion in the intron of GSVIVG01008260001 that is completely absent in the Merlot WT samples (Appendix S8). We propose that this genetic variant may be the causal mutation for the WB bud sport in the Merlot WB case investigated.

## DISCUSSION

### Developmental defects in the WB bud sport***—***

Our phenotypic measurements of the Dakapo and Merlot WB bud sports revealed new aspects of the WB phenotype that had previously been unknown. The most striking finding being how different the two instances of WB studied are from one another, with the Dakapo WB shoots having much smaller features in comparison the Merlot WB (Figs. 2A, B and 3E). Analysis of lateral meristem fate, leaf shape, and dormant buds further enforced how distinct the two instances of WB are (Figs. 2C, 4, and 5B, D). However, both WB cases were significantly smaller than their wild-type counterparts in every trait measured. The WB phenotype also seems to include development defects that have not been previously identified, such as subtle changes in leaf shape in both varieties (Fig. 4). The phenotypic measurements across the shoots of the WB bud sports show that not only are they smaller than their WT counterparts, but they also have defects in regulating overall shoot and leaf development. Our leaf size and shape data both seem to support that the WB leaves specifically seem to have very distinct developmental trajectories, with a) WB leaf areas not following the negative quadratic trend we expect to see as leaves age (Appendix S9) and b) WB leaves across the shoots having juvenile characteristics, such as deeper sinuses (Bryson et al., 2020) (Fig. 4). Identifying lateral meristem fates and analyzing internal bud morphologies also clarified developmental defects within the two instances of WB. These results suggested that the WB phenotype may be largely influenced by issues early on in meristem development, leading to a diverse array of developmental defects.

### Investigating the genetic basis of the WB bud sport***—***

Given the phenotypic differences between Dakapo WB and Merlot WB, it is possible that there are multiple genetic means of causing what is colloquially termed a “WB bud sport”. Mutational witch’s brooms are poorly described in angiosperms, although they are described from conifers (Zhuk et al. 2015), leading to few likely candidate genes in which mutations may drive the WB phenotype. Due to the large differences in phenotype between the two varieties, as well as none of the shared genes impacted by variants being good candidates for the bud sport, we propose that two different genetic variants cause the WB bud sport in the Dakapo and Merlot cases we investigated.

In Merlot, we identified a putative candidate gene for WB: GSVIVG01008260001. It is highly expressed in most tissue types, including buds, leaves, inflorescences, and roots (Fasoli et al. 2012), making it a promising candidate for a mutation with pleiotropic effects. GSVIVG01008260001 is orthologous to the gene AT1G49040 in *Arabidopsis*, which encodes STOMATAL CYTOKINESIS-DEFECTIVE 1 (AtSCD1). AtSCD1 is involved in the cytokinesis of guard mother cells and other leaf epidermal cells. However, AtSCD1 also appears to play a role in overall plant growth and development. In *Arabidopsis*, *scd1* mutants are smaller than WT plants, have reduced leaf expansion, and defects in flower morphology. The floral buds in *scd1* are smaller than WT due to early abortion in development and are highly branched as well (Falbel et al., 2003). The phenotype of the *scd1* floral buds is similar to the WB phenotype of Merlot WB buds, which are smaller than WT and also highly branching (Fig. 5D). The dwarfness and small leaves of *scd1* also match what we see in Merlot WB shoots. The abundant similarities between *scd1* mutants in *Arabidopsis* and the Merlot WB bud sport make GSVIVG01008260001 a strong candidate for one casual gene of the WB bud sport. Additionally, no other genes overlapping with SNPs or SVs unique to Merlot WB appear to be strong candidates. Most other genes identified as uniquely impacted by variants in Merlot WB do not appear to be involved in plant growth and development and/or are not truly genetically distinct in Merlot WB. Between the genetic evidence in the Merlot WB grapevine plants and phenotypic similarity to the *Arabidopsis* ortholog (Falbel et al., 2003), we propose GSVIVG01008260001 as a candidate causal gene for the WB bud sport in grapevine.

While we were able to identify a strong candidate in Merlot WB, no strong candidates were identified in the Dakapo WB. There are a few complicating factors that contributed to the difficulty of identifying a causal WB candidate in our Dakapo WB sample. For one, grapevine is highly heterozygous, which made it challenging to both accurately call and genotype SNPs and SVs within our samples. In addition, genetic chimeric variability, in which one cell layer has distinct genetic variants in comparison to the other cell layer, has repeatedly been identified in grapevine (Riaz et al. 2002; Franks et al. 2002). The phenotypic manifestation of a chimeric genetic variant depends on the cell layer(s) it is present within (Frank and Chitwood 2016). As a result, it is possible that the WB causal variant could be present in both WT and WB sequencing data, but present in distinct cell layer(s) between WT and WB. If the WB causal variant is chimeric in nature, it may not have been identified through our sequencing and variant calling. Finally, it is also possible that the WB bud sport could be the result of an epiallele as well, as was found with the mantled somaclonal variant that arises frequently in oil palm (Ong-Abdullah et al. 2015).

Ultimately, genetic transformation is necessary to prove the causal gene(s) of the WB bud sport. However, it is likely that an inducible mutant will need to be used to circumvent possible lethality due to issues that the bud sports have with rooting. As a result, the natural instances of the WB bud sport studied here provide invaluable natural mutants for studying whole plant development in grapevines. It is possible that the WB bud sport provides insight into developmental defects and interactions between developmental processes that might otherwise be impossible to study due to the inability of the WB bud sports to properly root and produce seed. Studying other occurrences of WB in the future will provide more insight into grapevine development and clarify the extent of the phenotypic and genetic diversity of “WB bud sports”.

### Somatic mutations in grapevine shoots and clones***—***

Our paired sequencing of WT and WB tissue from two instances of WB in grapevine also provided insight into somatic mutations both within plants and between clones. All samples had relatively similar counts of sample-unique SNPs when compared within variety (Table 1). We found between 410,608-411,425 of clone-specific SNPs in Dakapo, which is somewhat lower than what was previously reported in Zinfandel grapevine clones (Vondras et al., 2019). However, Vondras et al. (2019) called SNPs in the Zinfandel grapevine clones using a haplotype-resolved Zinfandel reference genome in order to identify clone-specific SNPs. This difference in methods likely accounts for most of the difference seen in clone-specific SNPs between our study and Vondras et al. (2019). Our data also provided insight into intra-organism mutations in grapevine, which to date and to our best knowledge have not been investigated. Our dataset revealed that the number of somatic mutations within one grapevine plant, when comparing distinct shoots (Merlot WT shoots and Merlot WB shoots), is similar to those found between grapevine clones (Table 1), with between 409,588-418,818 shoot-specific SNPs being identified in Merlot. The counts of shoot-specific SNPs in Merlot is somewhat higher than what has been identified in the past (4,973 SNPs in *Zostera marina* and 44,000-152,000 SNPs in *Populus trichocarpa*) (Hofmeister et al., 2020; Yu et al., 2020). However, the grapevine genome is much more heterozygous than that of *Z. marina* and *P. trichocarpa*, which likely lead to the identification of more false positive novel somatic mutations in our dataset. In addition, more novel somatic mutations would be expected in grapevine shoots given the high genetic divergence typically seen between clones.

Our long-read sequencing also provided insight into SV somatic mutations, which are relatively understudied in comparison to SNP somatic mutations, especially at the intra-organism level. We identified between 157-223 clonal-specific SVs in Dakapo, and between 102-224 shoot-specific SVs in Merlot. These findings align with our SNP data and support that the number of intra-organism somatic mutations in grapevine is similar to the number of inter-clone somatic mutations. The actual number of clonal-and shoot-specific SNPs and SVs is likely much lower than what was reported due to sequencing errors, alignment errors, etc. Regardless, these data provide insight into the accumulation of mutations within grapevine and supports the notion that grapevine clonal genetic diversity begins through novel somatic mutation accumulations on grapevine shoots, which are then clonally propagated.

## CONCLUSIONS

The WB bud sport provides a natural mutant in which to study developmental defects that might otherwise be impossible to study. Grapevine development is vastly different from that in *Arabidopsis*, and understanding this process and the genetic pathways involved will be invaluable in not only other perennial crop systems, but also in understanding liana development. However, studying the genes involved in grapevine development is difficult due to both traditional breeding and genetic transformation being relatively challenging and time consuming (Campos et al., 2021). Investigating the phenotypic defects and potential genetic basis of the WB bud sport has provided insight in grapevine development from buds to shoots. Future work in WB plants, especially with instances of the bud sport in new varieties and genetic backgrounds, will help deepen our understanding of development in grapevine, as well as other lianas and perennial crops.

## Supporting information

Appendix S1. Average leaf blade length and blade width at distinct nodes on the shoots for each sample

Appendix S2. Eigenleaves from the PCA comparing leaf shape between scaled Dakapo WT and Dakapo WB leaves, for PC 1-4

Appendix S3. Eigenleaves from the PCA comparing leaf shape between scaled Merlot WT and Merlot WB leaves, for PC 1-4

Appendix S4. Buds imaged using a dissecting microscope for all samples

Appendix S5. Number of SNPs with predicted SNP effects for all four samples individually

Appendix S6. SV types for all four samples individually

Appendix S7. Genetic candidates for casual variants of Witch's Broom in Dakapo and Merlot

Appendix S8. A diagram of the gene GSVIVG01008260001, the grapevine ortholog for AtSCD1

Appendix S9. The developmental trajectories of leaf area across shoots for all samples

## ACKNOWLEDGEMENTS

The authors thank Ileana Katzman, Erin Logan, and Brianna Wieferich for collecting and preparing grapevine herbarium specimens in 2022. We also thank the Genomics Core at Michigan State University and the Institute for Cyber-Enabled Research at Michigan State University for their services. This work is supported by Michigan State University, the USDA National Institute of Food and Agriculture MICL02572, and the NSF Plant Genome Research Program awards IOS-2310355, IOS-2310356, and IOS-2310357. EJR was supported by the University Distinguished Fellowship at Michigan State University.

## DATA AVAILABILITY

The phenotypic data from this study are available from the Dryad Digital Repository: X. Sequencing data from this study are provided on the NCBI Sequence Read Archive under BioProject PRJNA1020818. The code used for data analysis in this study is available on GitHub: https://github.com/eleanore-ritter/witchs-broom/.

## AUTHOR CONTRIBUTIONS

CN, EJR, and PC conceptualized and designed the study. PC provided plant material and collected shoot phenotype data. AK and EJR performed leaf landmarking. MQ performed CT scanning of buds. DHC, EJR, and MQ analyzed internal bud morphologies. SKKR isolated DNA and generated libraries for Illumina sequencing. AK and EJR validated potential candidate variants. EJR analyzed the data. CN, DHC, EJR, and PC assisted in data interpretation. EJR wrote the first draft of the manuscript. All authors assisted with final drafts of the manuscript.

## SUPPORTING INFORMATION

Additional supporting information may be found online in the Supporting Information section at the end of the article.

**Appendix S1.** Average leaf (A) blade length and (B) blade width at distinct nodes on the shoots for each sample, collected from 10 shoots each.

**Appendix S2.** Eigenleaves from the PCA comparing leaf shape between scaled Dakapo WT and Dakapo WB leaves, for PC 1-4.

**Appendix S3.** Eigenleaves from the PCA comparing leaf shape between scaled Merlot WT and Merlot WB leaves, for PC 1-4.

**Appendix S4.** Buds imaged using a dissecting microscope for (A) Dakapo WT, (B) Dakapo WB, (C) Merlot WT, and (D) Merlot WB samples. The vascular tissue projecting out of the Dakapo WB sample is directly right of the solid star symbol. The scale bars are 1 mm wide.

**Appendix S5.** Number of SNPs with predicted SNP effects for all four samples individually, when called against the 12X.v2 grapevine reference genome (Canaguier et al., 2017) using Illumina sequencing data.

**Appendix S6.** SV types for all four samples individually, when called against the 12X.v2 grapevine reference genome (Canaguier et al., 2017) using long-read sequencing data.

**Appendix S7.** Genetic candidates for casual variants of Witch’s Broom in Dakapo and Merlot.

**Appendix S8.** A diagram of the gene GSVIVG01008260001, the grapevine ortholog for AtSCD1. Exons are represented by black boxes along the gene body. The location and relative size of the 3.6 Kbp insertion present in Merlot WB is shown by the light purple line and triangle.

**Appendix S9.** The developmental trajectories of leaf area across shoots for (A) Dakapo WT, (B) Dakapo WB, (C) Merlot WT, and (D) Merlot WB. The blue line represents the linear model of the formula *y ∼ x + x^2^*, in which *y* is leaf area and *x* is node position. There was significant support for this negative quadratic relationship between leaf area and node in both Dakapo WT and Merlot WT (p<0.05 for both *x* and *x^2^* components for both varieties). However, there is not significant support for a negative quadratic relationship between leaf area and node in both Dakapo WB and Merlot WB (p>0.05 for both *x* and *x^2^* components for both varieties).

## LITERATURE CITED

1. Abramoff, P. J. Magalhães, and S. J. Ram. 2004. Image processing with ImageJ. Biophotonics international 11: 36–42.

2. Andrews, S. 2010. FastQC: a quality control tool for high throughput sequence data. http://www.bioinformatics.babraham.ac.uk/projects/fastqc.

3. Bettiga, L.J. 2013. Grape Pest Management, 3rd ed. University of California - Agriculture and Natural Resources, Oakland, California, USA.

4. Bolser, D., D. M. Staines, E. Pritchard, and P. Kersey. 2016. Ensembl Plants: Integrating Tools for Visualizing, Mining, and Analyzing Plant Genomics Data. Methods in molecular biology 1374: 115–140.

5. Broad Institute. 2019. Picard Toolkit, version 2.15.0. Broad Institute. Website: https://broadinstitute.github.io/picard/ [accessed 20 September 2023].

6. Bryson, A. E., M. Wilson Brown, J. Mullins, W. Dong, K. Bahmani, N. Bornowski, C. Chiu, et al. 2020. Composite modeling of leaf shape along shoots discriminates *Vitis* species better than individual leaves. Applications in plant sciences 8: e11404.

7. Buchfink, B., C. Xie, and D. H. Huson. 2015. Fast and sensitive protein alignment using DIAMOND. Nature methods 12: 59–60.

8. Campitelli, E. 2023. ggnewscale: Multiple Fill and Colour Scales in ’ggplot2’, version 0.4.8. Website: https://CRAN.R-project.org/package=ggnewscale [accessed 20 September 2023].

9. Campos, G., C. Chialva, S. Miras, and D. Lijavetzky. 2021. New Technologies and Strategies for Grapevine Breeding Through Genetic Transformation. Frontiers in plant science 12: 767522.

10. Canaguier, A., J. Grimplet, G. Di Gaspero, S. Scalabrin, E. Duchêne, N. Choisne, N. Mohellibi, et al. 2017. A new version of the grapevine reference genome assembly (12X.v2) and of its annotation (VCost.v3). Genomics data 14: 56–62.

11. Carbonell-Bejerano, P., C. Royo, R. Torres-Pérez, J. Grimplet, L. Fernandez, J. M. Franco-Zorrilla, D. Lijavetzky, et al. 2017. Catastrophic Unbalanced Genome Rearrangements Cause Somatic Loss of Berry Color in Grapevine. Plant physiology 175: 786–801.

12. Cheng, C., C. Jiao, S. D. Singer, M. Gao, X. Xu, Y. Zhou, Z. Li, et al. 2015. Gibberellin-induced changes in the transcriptome of grapevine (*Vitis labrusca* × *V. vinifera*) cv. Kyoho flowers. BMC genomics 16: 128.

13. Cheng, C.-Y., V. Krishnakumar, A. P. Chan, F. Thibaud-Nissen, S. Schobel, and C. D. Town. 2017. Araport11: a complete reannotation of the *Arabidopsis thaliana* reference genome. The Plant journal: for cell and molecular biology 89: 789–804.

14. Chitwood, D. H., J. Mullins, Z. Migicovsky, M. Frank, R. VanBuren, and J. P. Londo. 2021. Vein-to-blade ratio is an allometric indicator of leaf size and plasticity. American journal of botany 108: 571–579.

15. Chitwood, D. H., S. M. Rundell, D. Y. Li, Q. L. Woodford, T. T. Yu, J. R. Lopez, D. Greenblatt, et al. 2016. Climate and Developmental Plasticity: Interannual Variability in Grapevine Leaf Morphology. Plant physiology 170: 1480–1491.

16. Cingolani, P., V. M. Patel, M. Coon, T. Nguyen, S. J. Land, D. M. Ruden, and X. Lu. 2012. Using *Drosophila melanogaster* as a Model for Genotoxic Chemical Mutational Studies with a New Program, SnpSift. Frontiers in genetics 3: 35.

17. Constantin, A. E., and I. Patil. 2021. ggsignif: R package for displaying significance brackets for ‘ggplot2.’ PsyArxiv.

18. Danecek, P., J. K. Bonfield, J. Liddle, J. Marshall, V. Ohan, M. O. Pollard, A. Whitwham, et al. 2021. Twelve years of SAMtools and BCFtools. GigaScience 10.

19. De Coster, W., S. D’Hert, D. T. Schultz, M. Cruts, and C. Van Broeckhoven. 2018. NanoPack: visualizing and processing long-read sequencing data. Bioinformatics 34: 2666–2669.

20. DePristo, M. A., E. Banks, R. Poplin, K. V. Garimella, J. R. Maguire, C. Hartl, A. A. Philippakis, et al. 2011. A framework for variation discovery and genotyping using next-generation DNA sequencing data. Nature genetics 43: 491–498.

21. Dryden, I. L., and K. V. Mardia. 2016. Statistical Shape Analysis: With Applications in R. John Wiley & Sons.

22. Durinck, S., P. T. Spellman, E. Birney, and W. Huber. 2009. Mapping identifiers for the integration of genomic datasets with the R/Bioconductor package biomaRt. Nature protocols 4: 1184– 1191.

23. Falbel, T. G., L. M. Koch, J. A. Nadeau, J. M. Segui-Simarro, F. D. Sack, and S. Y. Bednarek. 2003. SCD1 is required for cytokinesis and polarized cell expansion in *Arabidopsis thaliana* [corrected]. Development 130: 4011–4024.

24. Foster, T. M., and M. J. Aranzana. 2018. Attention sports fans! The far-reaching contributions of bud sport mutants to horticulture and plant biology. Horticulture research 5: 44.

25. Frank, M. H., and D. H. Chitwood. 2016. Plant chimeras: The good, the bad, and the ‘Bizzaria’. Developmental biology 419: 41–53.

26. Franks, T., R. Botta, M. R. Thomas, and J. Franks. 2002. Chimerism in grapevines: implications for cultivar identity, ancestry and genetic improvement. Theoretical and applied genetics 104: 192–199.

27. Gerrath, J., U. Posluszny, and L. Melville. 2015. Taming the Wild Grape: Botany and Horticulture in the Vitaceae. Springer.

28. Hofmeister, B. T., J. Denkena, M. Colomé-Tatché, Y. Shahryary, R. Hazarika, J. Grimwood, S. Mamidi, et al. 2020. A genome assembly and the somatic genetic and epigenetic mutation rate in a wild long-lived perennial *Populus trichocarpa*. Genome biology 21: 259.

29. Jaillon, O., J.-M. Aury, B. Noel, A. Policriti, C. Clepet, A. Casagrande, N. Choisne, et al. 2007. The grapevine genome sequence suggests ancestral hexaploidization in major angiosperm phyla. Nature 449: 463–467.

30. Jeffares, D. C., C. Jolly, M. Hoti, D. Speed, L. Shaw, C. Rallis, F. Balloux, et al. 2017. Transient structural variations have strong effects on quantitative traits and reproductive isolation in fission yeast. Nature communications 8: 14061.

31. Jung, H. Y. 2002. ‘*Candidatus* Phytoplasma castaneae’, a novel phytoplasma taxon associated with chestnut witches’ broom disease. International journal of systematic and evolutionary microbiology 52: 1543–1549.

32. Khan, A. J., S. Botti, A. M. Al-Subhi, D. E. Gundersen-Rindal, and A. F. Bertaccini. 2002. Molecular identification of a new phytoplasma associated with alfalfa witches’-broom in oman. Phytopathology 92: 1038–1047.

33. Li, H. 2021. New strategies to improve minimap2 alignment accuracy. Bioinformatics 37: 4572– 4574.

34. Li, H., and R. Durbin. 2009. Fast and accurate short read alignment with Burrows–Wheeler transform. Bioinformatics 25: 1754–1760.

35. Li, H., B. Handsaker, A. Wysoker, T. Fennell, J. Ruan, N. Homer, G. Marth, et al. 2009. The Sequence Alignment/Map format and SAMtools. Bioinformatics 25: 2078–2079.

36. Martin, M. 2011. Cutadapt removes adapter sequences from high-throughput sequencing reads. EMBnet.journal 17: 10–12.

37. McKenna, A., M. Hanna, E. Banks, A. Sivachenko, K. Cibulskis, A. Kernytsky, K. Garimella, et al. 2010. The Genome Analysis Toolkit: a MapReduce framework for analyzing next-generation DNA sequencing data. Genome research 20: 1297–1303.

38. Montano, H. G., R. E. Davis, E. L. Dally, S. Hogenhout, P. Pimentel, and P. S. T. Brioso. 2001. ’ *Candidatus* Phytoplasma brasiliense ‘, a new phytoplasma taxon associated with hibiscus witches’ broom disease. Website https://pubag.nal.usda.gov/download/18621/pdf [accessed 12 April 2023].

39. Ong-Abdullah, M., J. M. Ordway, N. Jiang, S.-E. Ooi, S.-Y. Kok, N. Sarpan, N. Azimi, et al. 2015. Loss of Karma transposon methylation underlies the mantled somaclonal variant of oil palm. Nature 525: 533–537.

40. Pacific Biosciences. 2021. pbsv, version 2.8.0. Website: https://github.com/PacificBiosciences/pbsv [accessed 20 September 2023].

41. Porebski, S., Bailey, L.G. and Baum, B.R. 1997. Modification of a CTAB DNA Extraction Protocol for Plants Containing High Polysaccharide and Polyphenol Components. Plant Molecular Biology Reporter 15: 8–15.

42. Qiagen. 2015. QIAGEN® Genomic DNA Handbook.

43. Quinlan, A. R., and I. M. Hall. 2010. BEDTools: a flexible suite of utilities for comparing genomic features. Bioinformatics 26: 841–842.

44. R Core Team. 2022. R: A Language and Environment for Statistical Computing. R Foundation for Statistical Computing. Website: https://www.R-project.org/ [accessed 20 September 2023].

45. Riaz, S., K. E. Garrison, G. S. Dangl, J.-M. Boursiquot, and C. P. Meredith. 2002. Genetic Divergence and Chimerism within Ancient Asexually Propagated Winegrape Cultivars. Journal of the American Society for Horticultural Science 127.

46. RStudio Team. 2022. RStudio: Integrated Development Environment for R. RStudio, PBC. Website: https://posit.co/products/open-source/rstudio/ [accessed 20 September 2023].

47. Sedlazeck, F. J., P. Rescheneder, M. Smolka, H. Fang, M. Nattestad, A. von Haeseler, and M. C. Schatz. 2018. Accurate detection of complex structural variations using single-molecule sequencing. Nature methods 15: 461–468.

48. Shumate, A., and S. L. Salzberg. 2021. Liftoff: accurate mapping of gene annotations. Bioinformatics 37: 1639–1643.

49. Srinivasan, C., and M. G. Mullins. 1981. Physiology of Flowering in the Grapevine — a Review. American journal of enology and viticulture 32: 47–63.

50. Urich MA, Nery JR, Lister R, Schmitz RJ, Ecker JR. 2015. MethylC-seq library preparation for base-resolution whole-genome bisulfite sequencing. Nature Protocols 10(3): 475–83.

51. Van der Auwera, G. A., M. O. Carneiro, C. Hartl, R. Poplin, G. Del Angel, A. Levy-Moonshine, T. Jordan, et al. 2013. From FastQ data to high confidence variant calls: the Genome Analysis Toolkit best practices pipeline. Current protocols in bioinformatics 11(1110): 11.10.1–11.10.33.

52. Vondras, A. M., A. Minio, B. Blanco-Ulate, R. Figueroa-Balderas, M. A. Penn, Y. Zhou, D. Seymour, et al. 2019. The genomic diversification of grapevine clones. BMC genomics 20: 972.

53. Wang, K., M. Li, and H. Hakonarson. 2010. ANNOVAR: functional annotation of genetic variants from high-throughput sequencing data. Nucleic acids research 38: e164.

54. Wickham, H. 2016. Programming with ggplot2. In H. Wickham [ed.], ggplot2: Elegant Graphics for Data Analysis, 241–253. Springer International Publishing, Cham.

55. Wick, R. R., L. M. Judd, C. L. Gorrie, and K. E. Holt. 2017. Completing bacterial genome assemblies with multiplex MinION sequencing. Microbial genomics 3: e000132.

56. Wilke, C.O. 2020. cowplot: Streamlined Plot Theme and Plot Annotations for ’ggplot2’, version 1.1.1. Website: https://CRAN.R-project.org/package=cowplot [accessed 20 September 2023].

57. Yu, L., C. Boström, S. Franzenburg, T. Bayer, T. Dagan, and T. B. H. Reusch. 2020. Somatic genetic drift and multilevel selection in a clonal seagrass. Nature ecology & evolution 4: 952–962.

58. Zhuk, E., G. Vasilyeva, and S. Goroshkevich. 2015. Witches’ broom and normal crown clones from the same trees of *Pinus sibirica*: a comparative morphological study. Trees 29: 1079– 1090.

