## Appendix S1. Average leaf blade length and blade width at distinct nodes on the shoots for each sample for "From buds to shoots: Insights into grapevine development from the Witch’s Broom bud sport"

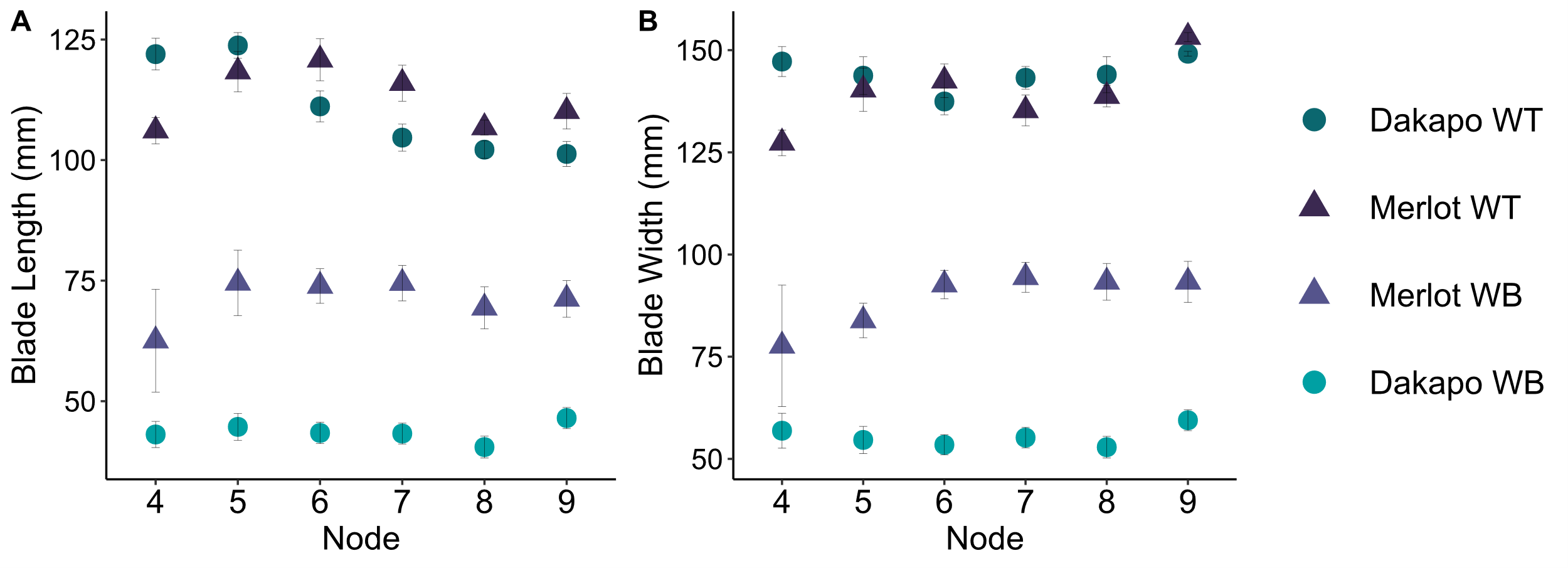

**Appendix S1.** Average leaf **(A)** blade length and **(B)** blade width at distinct nodes on the shoots for each sample, collected from 10 shoots each.
