## Appendix S2. Eigenleaves from the PCA comparing leaf shape between scaled Dakapo WT and Dakapo WB leaves, for PC 1-4 for "From buds to shoots: Insights into grapevine development from the Witch’s Broom bud sport"

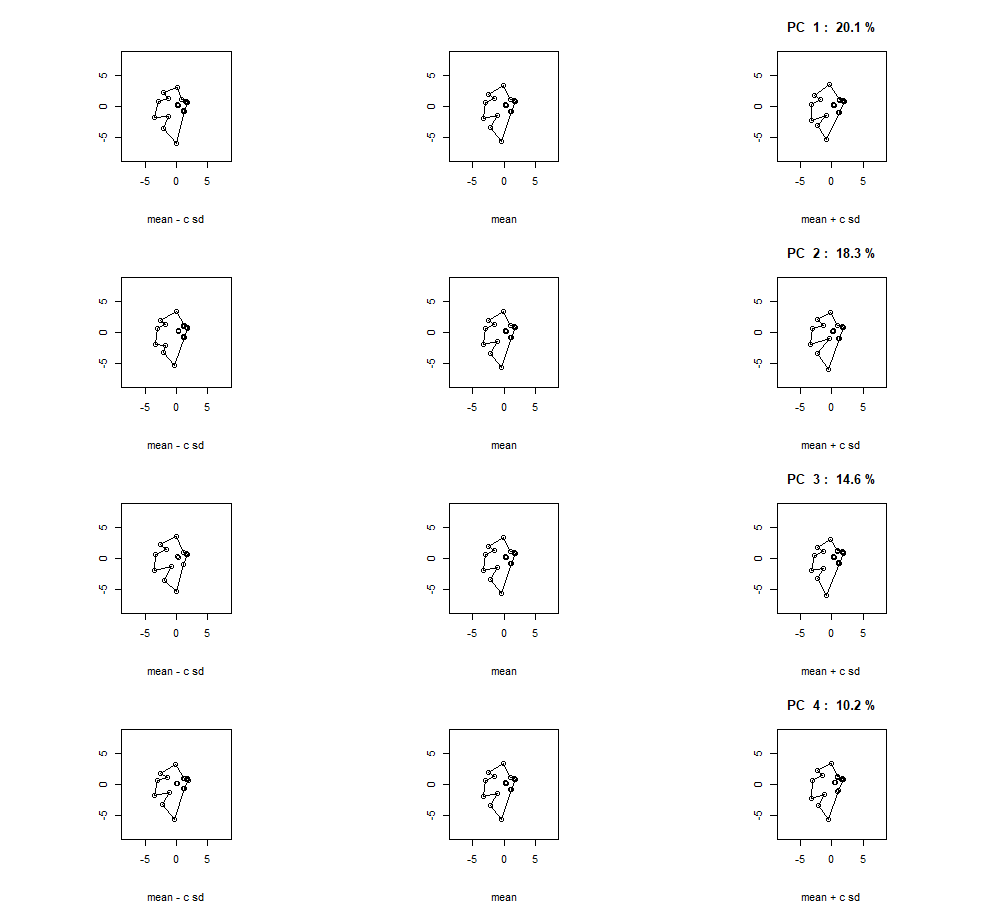

**Appendix S2.** Eigenleaves from the PCA comparing leaf shape between scaled Dakapo WT and Dakapo WB leaves, for PC 1-4.
