## Appendix S3. Eigenleaves from the PCA comparing leaf shape between scaled Merlot WT and Merlot WB leaves, for PC 1-4 for "From buds to shoots: Insights into grapevine development from the Witch’s Broom bud sport"

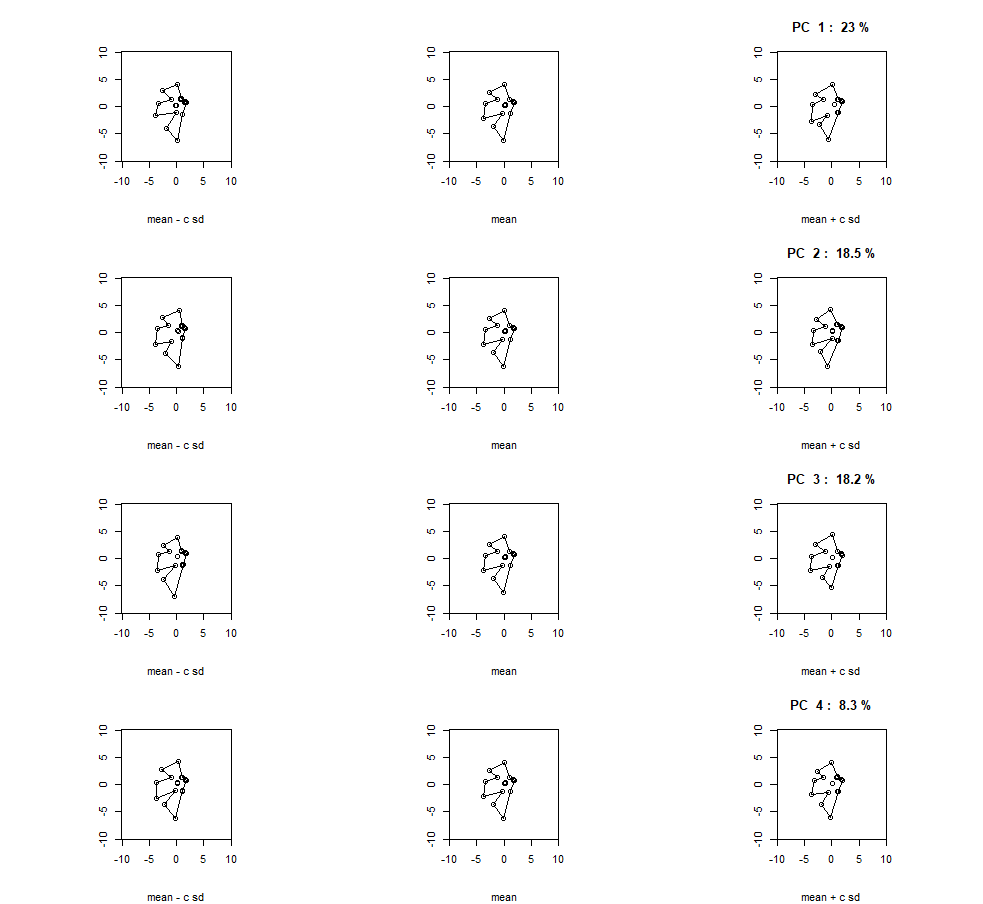

**Appendix S3.** Eigenleaves from the PCA comparing leaf shape between scaled Merlot WT and Merlot WB leaves, for PC 1-4.
