## Appendix S4. Buds imaged using a dissecting microscope for all samples for "From buds to shoots: Insights into grapevine development from the Witch’s Broom bud sport"

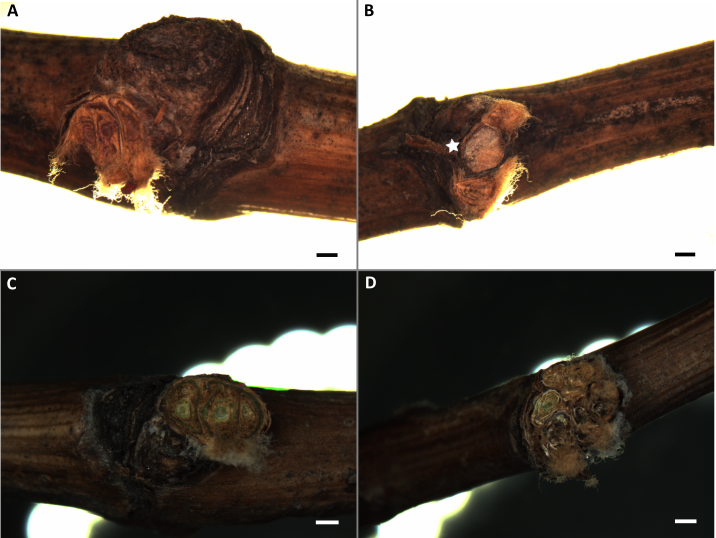

**Appendix S4.** Buds imaged using a dissecting microscope for **(A)** Dakapo WT, **(B)** Dakapo WB, **(C)** Merlot WT, and **(D)** Merlot WB samples. The vascular tissue projecting out of the Dakapo WB sample is directly right of the solid star symbol. The scale bars are 1 mm wide.
