## Appendix S5. Number of SNPs with predicted SNP effects for all four samples individually for "From buds to shoots: Insights into grapevine development from the Witch’s Broom bud sport"

|  | **Dakapo** | | **Merlot** | |
| --- | --- | --- | --- | --- |
| **Predicted SNP Effects** | **WT** | **WB** | **WT** | **WB** |
| **downstream** | 302954 | 302701 | 308529 | 309713 |
| **intergenic** | 4485234 | 4494808 | 4559676 | 4570032 |
| **intronic** | 2451458 | 2454637 | 2594527 | 2597389 |
| **ncRNA_exonic** | 17 | 22 | 14 | 14 |
| **splicing** | 2448 | 2428 | 2456 | 2459 |
| **upstream** | 369709 | 372057 | 377116 | 377983 |
| **upstream; downstream** | 40420 | 40459 | 41191 | 41112 |
| **UTR3** | 68803 | 68712 | 70248 | 70432 |
| **UTR5** | 47161 | 47189 | 48194 | 48313 |
| **exonic** |  |  |  |  |
| frameshift | 7827 | 7738 | 7814 | 7798 |
| nonframeshift | 3446 | 3443 | 3515 | 3532 |
| nonsynonymous | 115910 | 115683 | 117985 | 117927 |
| stop gain | 3382 | 3404 | 3391 | 3389 |
| stop loss | 479 | 485 | 495 | 499 |
| synonymous | 90373 | 89946 | 92738 | 92715 |
| unknown | 148 | 145 | 147 | 162 |

**Appendix S5.** Number of SNPs with predicted SNP effects for all four samples individually, when called against the 12X.v2 grapevine reference genome (Canaguier et al., 2017) using Illumina sequencing data.
