## Appendix S6. SV types for all four samples individually for "From buds to shoots: Insights into grapevine development from the Witch’s Broom bud sport"

|  | **Dakapo** | | **Merlot** | |
| --- | --- | --- | --- | --- |
|  | **WT** | **WB** | **WT** | **WB** |
| Deletions | 27,173 | 27,420 | 28,495 | 28,122 |
| Insertions | 24,492 | 24,911 | 25,677 | 25,326 |
| Inversions | 65 | 71 | 67 | 63 |
| Transversions | 662 | 754 | 636 | 575 |
| Duplications | 57 | 58 | 56 | 52 |

**Appendix S6.** SV types for all four samples individually, when called against the 12X.v2 grapevine reference genome (Canaguier et al., 2017) using long-read sequencing data.
