## Appendix S8. A diagram of the gene GSVIVG01008260001, the grapevine ortholog for AtSCD1 for "From buds to shoots: Insights into grapevine development from the Witch’s Broom bud sport"

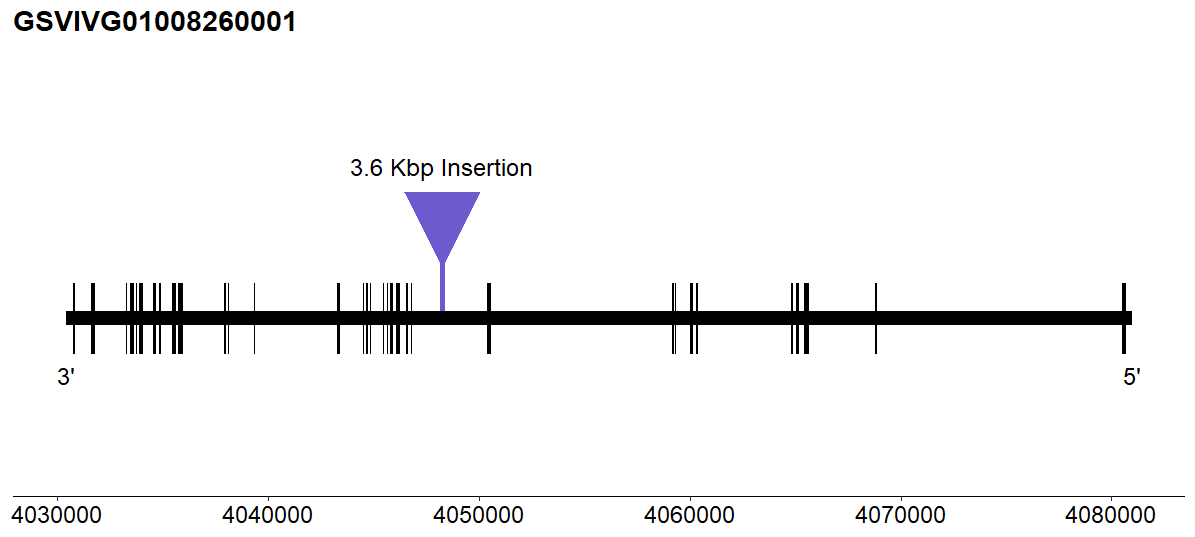

**Appendix S8.** A diagram of the gene GSVIVG01008260001, the grapevine ortholog for AtSCD1. Exons are represented by black boxes along the gene body. The location and relative size of the 3.6 Kbp insertion present in Merlot WB is shown by the light purple line and triangle.
