## Appendix S9. The developmental trajectories of leaf area across shoots for all samples for "From buds to shoots: Insights into grapevine development from the Witch’s Broom bud sport"

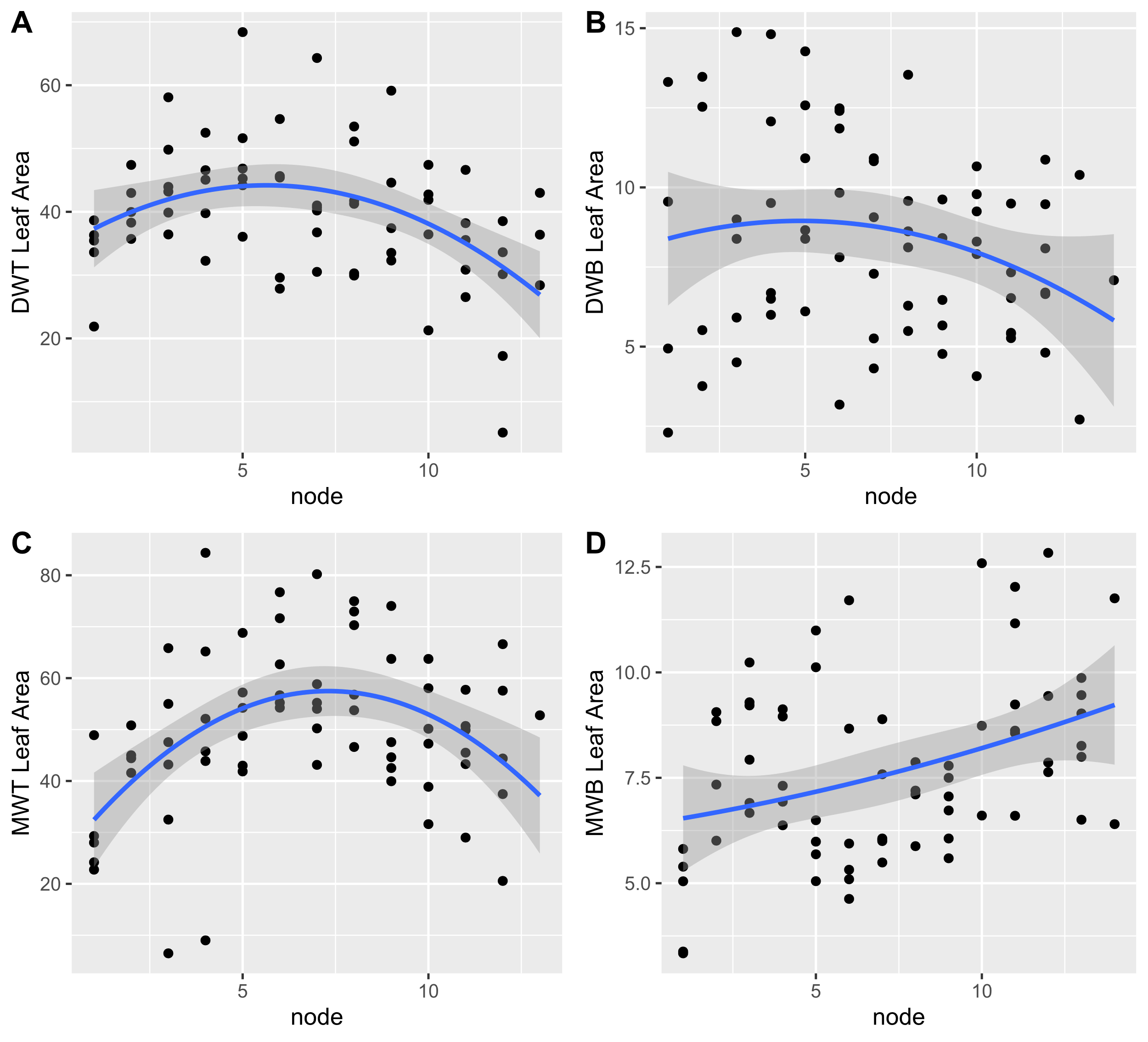

**Appendix S9.** The developmental trajectories of leaf area across shoots for **(A)** Dakapo WT, **(B)** Dakapo WB, **(C)** Merlot WT, and **(D)** Merlot WB. The blue line represents the linear model of the formula *y ~ x + x^2^*, in which *y* is leaf area and *x* is node position. There was significant support for this negative quadratic relationship between leaf area and node in both Dakapo WT and Merlot WT (p<0.05 for both *x* and *x^2^* components for both varieties).  However, there is not significant support for a negative quadratic relationship between leaf area and node in both Dakapo WB and Merlot WB (p>0.05 for both *x* and *x^2^* components for both varieties).
